# Rapid “recycling” of logical algorithm representations in fronto-parietal reasoning systems following computer programming instructions

**DOI:** 10.1101/2025.01.08.631982

**Authors:** Yun-Fei Liu, Marina Bedny

**Affiliations:** Johns Hopkins University

## Abstract

Programming is a cornerstone of modern society, yet its cognitive and neural basis remains poorly understood. In this study, we test the hypothesis that programming “recycles” pre-existing neural mechanisms and representations in fronto-parietal reasoning networks. Using fMRI, we scanned programming-naïve undergraduates (n=22) before (PRE) and after (POST) an introductory Python course. During the PRE scan, participants viewed pseudocode (plain English descriptions of algorithms), and during the POST scan, they read Python code. We found that a left-lateralized fronto-parietal network, previously implicated in programming experts, distinguished between “for” loops and “if” conditionals across both pseudocode and Python code. Representational similarity analysis revealed consistent representations of algorithms across formats (code/pseudocode) and learning stages. Furthermore, such representations encode abstract meanings rather than superficial features. Our findings demonstrate that programming not only recycles pre-existing neural resources evolved for logical reasoning, but the recycling takes place rapidly with only a single semester of training.

## Introduction

Modern life depends heavily on computer programming and its many applications, the latest of which is artificial intelligence. Yet, little is known about how the human brain enables this essential cultural skill. The neural recycling hypothesis proposes that cultural abilities are acquired by reusing preexisting neural representations engaged by related cognitive functions (Dehaene & Cohen, 2007). For example, literacy is thought to modify visual object representations in the lateral ventral visual stream and their connectivity with language networks (Dehaene-Lambertz, Monzalvo, & Dehaene, 2018; Dehaene, Cohen, Morais, & Kolinsky, 2015; X. Feng, Monzalvo, Dehaene, & Dehaene-Lambertz, 2022; McCandliss, Cohen, & Dehaene, 2003). Mathematics is hypothesized to recycle approximate number representations in parietal cortices (Brannon & Merritt, 2011; Cantlon & Brannon, 2007; Feigenson, Dehaene, & Spelke, 2004; Nieder, 2021; Tudusciuc & Nieder, 2009).

The study of programming offers a unique opportunity to test the neural recycling framework. Unlike reading and math, which are often learned in childhood, programming is typically first encountered in adolescence or adulthood, even in industrialized societies. This makes it possible to acquire high-resolution data from the same individuals both before and after their initial exposure to programming to ask whether programming reuses neural representations that predate programming literacy.

Although the ability to understand and write code depends on a number of cognitive abilities, one essential element of coding is the representation and manipulation of logical algorithms, such as “for” loops and “if” conditionals (Ivanova et al., 2020; Liu, Kim, Wilson, & Bedny, 2020). One possibility is that such code-relevant algorithm representations develop as a result of programming instruction and practice. In the current study we tested the alternative neural recycling hypothesis: that code-relevant algorithm representations are present in fronto-parietal reasoning system even prior to first exposure to coding and are “recycled” by programming instruction.

Several sources of evidence suggest that fronto-parietal reasoning systems play a role in programming. First, behavioral studies of individual differences find that reasoning abilities prior to instruction predict coding performance (Anderson, Farrell, & Sauers, 1984; Farghaly & El-Kafrawy, 2021; Graafsma et al., 2023; McCoy & Burton, 1988; Prat, Madhyastha, Mottarella, & Kuo, 2020; Shute, 1991). Recent evidence also suggests that that expert programmers recruit fronto-parietal reasoning networks during code comprehension and generation (Castelhano et al., 2018; Endres, Karas, Hu, Kovelman, & Weimer, 2021; Floyd, Santander, & Weimer, 2017; Hishikawa, Yoshinaga, Togo, Hongo, & Hanakawa, 2023; Ikutani et al., 2021; Ivanova et al., 2020; Krueger et al., 2020; Liu et al., 2020; Siegmund et al., 2014; Xu, Li, & Liu, 2021). These networks show higher responses during code comprehension than during difficulty-matched memory tasks. Moreover, activity patterns in these networks distinguish between code-relevant algorithms, including “for” loops and “if” conditionals (Ikutani et al., 2021; Liu et al., 2020; Srikant, Lipkin, Ivanova, Fedorenko, & O’Reilly, 2022). In the current study, we asked whether fronto-parietal reasoning networks represent this algorithmic distinction even prior to coding instruction.

We measured the functional properties of neural networks supporting Python code comprehension before and after a group of Johns Hopkins undergraduates (n = 22) took their first programming course (Gateway Computing: Python). A key challenge in this endeavor is measuring programming-relevant representations in people who do not yet know the surface form—so-called “syntax”—of programming scripts. Presenting naïve participants with actual code is problematic, as any lack of response is likely related to their inability to understand the surface form rather than the meaning of the stimuli. To address this, we designed custom “pseudocode” passages that describe programming algorithms in plain English (Figure 1).

**Figure 1.**
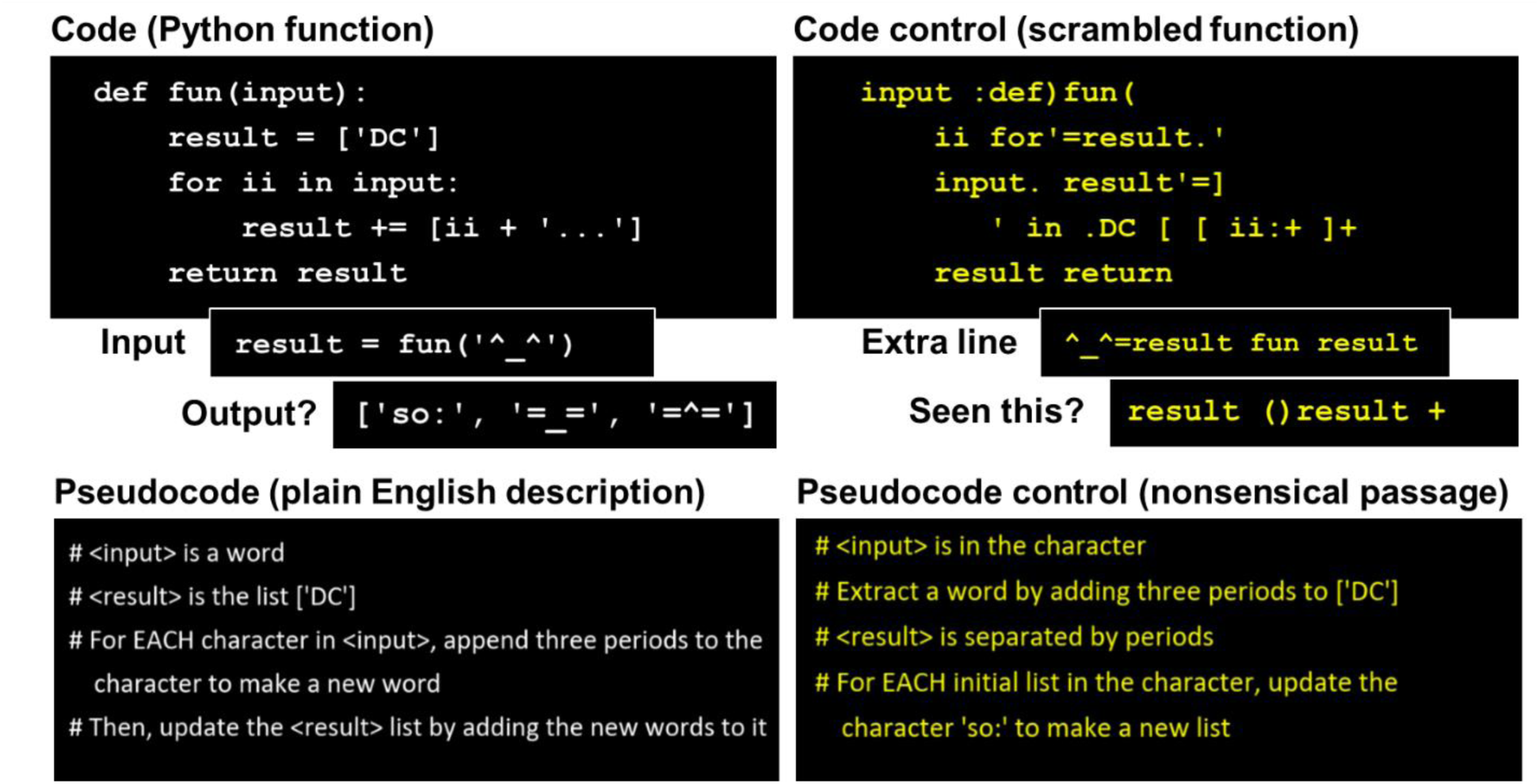
Example stimuli and task design. The example stimuli were presented to participants during the “reading” phase of a trial (20s). During the “input” phase (6s), the stimuli remained on the screen with an extra line. For pseudocode reading trials, the extra line was in the format “Now <INPUT> is the word ‘_____’”. For pseudocode control trials, the extra line was a sentence. During the “question” phase (4s), a single line was presented. For code or pseudocode reading trials, participants answered whether the line indicated the correct output of the algorithm. For control trials, participants answered whether the line was present during the “input” phase. Each trial began with a 0.5s fixation cross. “Input” and “question” phases were also separated by a 0.5s fixation cross. Inter-trial interval was 5s.

Participants viewed pseudocode passages describing “for” and “if” programming snippets before taking their first Python course (the PRE scan). They then returned for a second scan after the semester and saw analogous “for” and “if” algorithms in the form of actual code snippets written in Python, as well as a new set of pseudocode passages (the POST scan). The same participants also completed a localizer task to identify and distinguish fronto-parietal logical reasoning networks from fronto-temporal language networks (Kanjlia, Lane, Feigenson, & Bedny, 2016; Monti, Parsons, & Osherson, 2009, 2012). Previous work with expert programmers found significantly more overlap with reasoning than language systems (Ivanova et al., 2020; Liu et al., 2020; Xu et al., 2021), and we tested for this signature in beginning learners. We also measured the lateralization of the coding network, as prior studies have shown left-lateralized responses to code in experts (Castelhano et al., 2018; Floyd et al., 2017; Ikutani et al., 2021; Ivanova et al., 2020; Liu et al., 2020; Siegmund et al., 2014; Xu et al., 2021).

## Results

The cultural recycling framework predicts that the neural networks supporting program comprehension emerge rapidly with instruction and resemble those observed in prior studies with experts. This framework also makes a strong prediction: neural networks that later support code comprehension after instruction should contain neural populations representing code-relevant algorithms even before instruction. To test this prediction, we functionally identified code-responsive fronto-parietal networks in each individual after instruction and then probed the representations of these same networks prior to learning using multivariate methods. Specifically, we employed decoding to test for the distinction between “for” and “if” algorithms in the fronto-parietal systems both before and after instruction. Additionally, we conducted representational similarity analysis (RSA) to compare fronto-parietal responses before and after instruction and to examine their representational content in detail, extending beyond the for/if algorithm distinction.

An alternative hypothesis posits that a consistent, expert-like neural network requires many years of instruction and practice to develop. Under this hypothesis, new learners might exhibit diffuse or variable responses. It is also possible that different networks, such as the language network, may initially support code comprehension; over time, these functions could transfer to fronto-parietal systems. Furthermore, the neural population representations of algorithms within fronto-parietal systems, which can be detected with multivariate techniques, may emerge only after extensive practice.

### Participants exhibited program comprehension during in-scanner tasks and additional behavioral assessments

Participants self-reported no prior programming experience before taking the introductory programming course. At the end of the semester, participants’ acquisition of programming knowledge was attested by their performance in the course. The average grade was a B but varied across students, with none of the participants failing the course (mean=86.8%, SD=8.9%, range: 61.6% - 96.1%). Additionally, we assessed participants’ learning outcome with a simple five-option multiple-choice test on their knowledge of Python “syntax” (accuracy mean=70%, SD=12.1%; random guess chance level=20%), as well as a difficult open-ended fill-in-the-blank test on their ability to apply programming knowledge to implement specified algorithms (accuracy mean=31.1%, SD=16.4%; chance level=0%).

In-scanner performance was above chance on all tasks during both the PRE and POST scans. Participants understood programming algorithms described in pseudocode even prior to instruction, as evidenced by their high task performance on the “is this output correct” task during the pre-instruction scan (task accuracy mean=84%, SD=5.9%; response time mean=1.73s, SD=0.26s). Performance during the POST scan was also high for both the pseudocode and code reading tasks (pseudocode accuracy mean=88.3%, SD=7.3%; r.t. mean=1.39s, SD=0.31s. code accuracy mean=84.4%, SD=8.3%; r.t. mean=1.42s, SD=0.31s. Supplementary Figure 1). The memory control tasks were more difficult than their corresponding pseudocode or code reading tasks in terms of behavioral performance. Please see supplementary results for the statistics.

### The fronto-parietal reasoning network responds to code before and after initial programming instruction: univariate analysis

After a single semester of Python instruction, a left-lateralized fronto-parietal network responded more to Python code than a memory control task (laterality index: mean= -.56, SD=0.35, signed-rank test against 0: W=11, p<0.001. Figure 2a. See Table 1 for activated clusters. Also see Supplementary Figure 2 for laterality index). This network is highly similar to what has previously been observed for expert programmers (Liu et al., 2020). As with experts, a cluster of activation was also observed the posterior aspect of the left middle temporal gyrus (Liu et al., 2020). The new learner’s Python comprehension network overlapped more with fronto-parietal circuits involved in logical reasoning than with the fronto-temporal language network (Figure 2b; overlap with logic measured in dice coefficients, mean=0.44, SD=0.14; overlap with language: mean=0.17, SD=0.07; logic vs. language overlap permutation tested against each other p<0.001).

**Figure 2.**
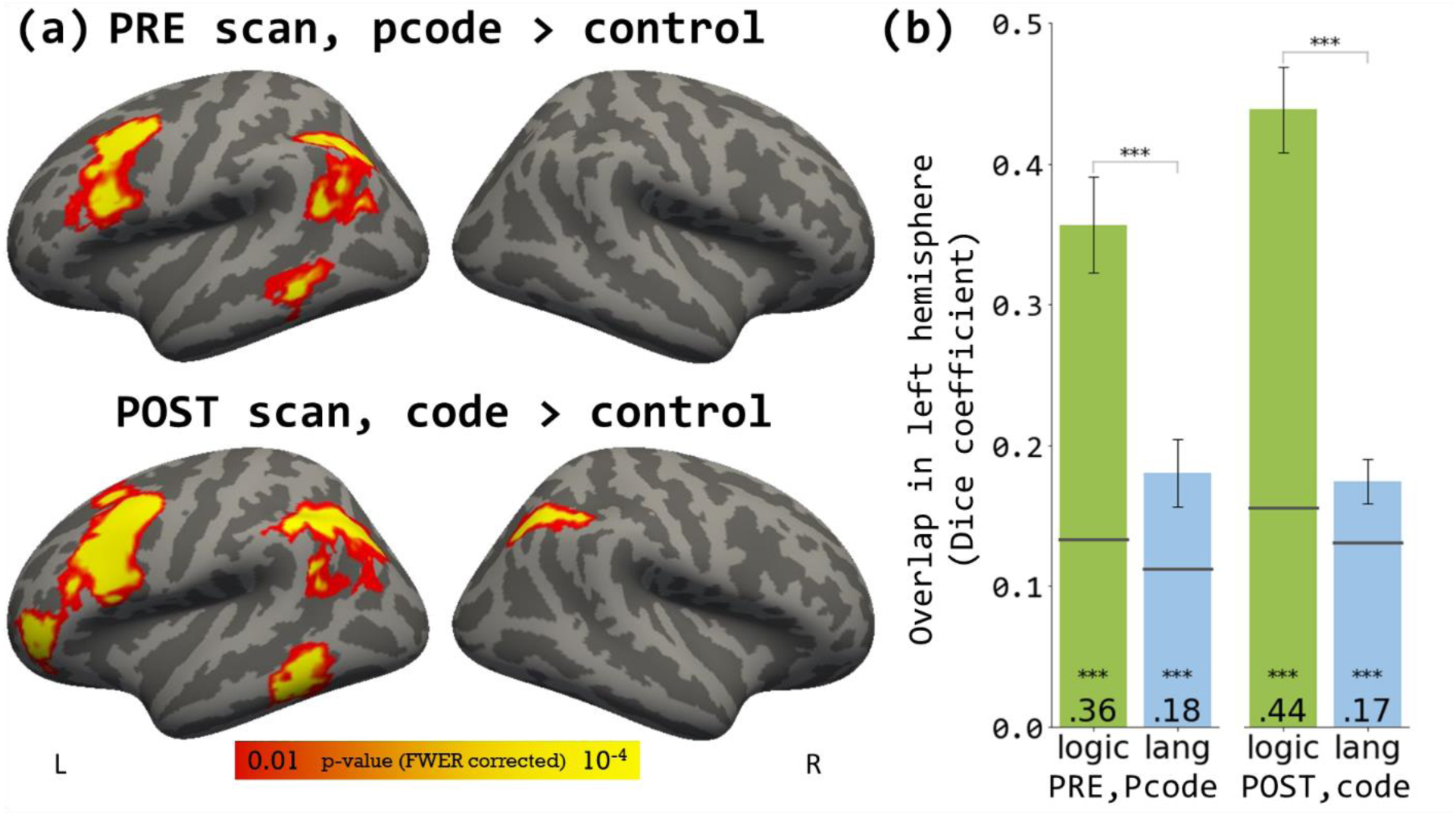
Univariate responses. (a) Top: Pseudocode (pcode) > memory control contrast during the PRE scan; bottom: Code > memory control contrast during the POST scan. Corrected for FWER with cluster-forming threshold of p<0.01 and cluster-wise p<0.05. (b) Overlap (Dice coefficient) between spatial activation patterns in (a) with logic or language (lang) networks localized within the same individuals. Error bars indicate standard error. Horizontal black lines on each bar indicate chance level overlap given the observed number of active vertices. Statistical significance of each bar was tested against respective chance level. ***p<0.001. For the activation maps with logic and language network overlaid, please see Supplementary Figure 3.

Even prior to programming instruction (i.e., during the PRE scan), a similar left-lateralized fronto-parietal network was already engaged during the comprehension of plain English pseudocode algorithms (compared to memorizing sentences containing the same words but without executable algorithms) (laterality index: mean=-0.61, SD=0.33, W=5, p<0.001). Unlike responses to real code, the activation did not extend into the frontal pole (Figure 2a. Table 1). Pseudocode also exhibited greater overlap with the logic than the language network, despite being presented in written English (overlap with logic: mean=0.36, SD=0.16, against chance p<0.001; with language: mean=0.18, SD=0.11, against chance p<0.001. Logic vs. language permutation test, p<0.001). Similar but weaker neural response to pseudocode were reinstated during the POST scan (see Supplementary results and Supplementary Figure 3).

### Neural populations in the reasoning network represent Python-relevant algorithms before and after programming instruction: “for” vs. “if” MVPA decoding

As previously shown for coding experts, in Python students with just one semester of experience, code-responsive intraparietal sulcus (IPS) and lateral prefrontal cortex (PFC) vertices showed sensitivity to the distinction between “for” loop and “if” conditional algorithms in Python code during the POST scan (Wilcoxon signed-rank test against chance: IPS W=218.0, p<0.001; PFC W=180.0, p<0.001. Control region primary auditory cortex A1 W=93, p=0.53. See Supplementary Results for mean and SD of decoding accuracy). In addition, decoding accuracy in the IPS and the PFC was both significantly greater than in the control region A1 (IPS vs A1: W=212.5, p<0.01. PFC vs A1: W=216.5, p<0.01. Figure 3a).

**Figure 3.**
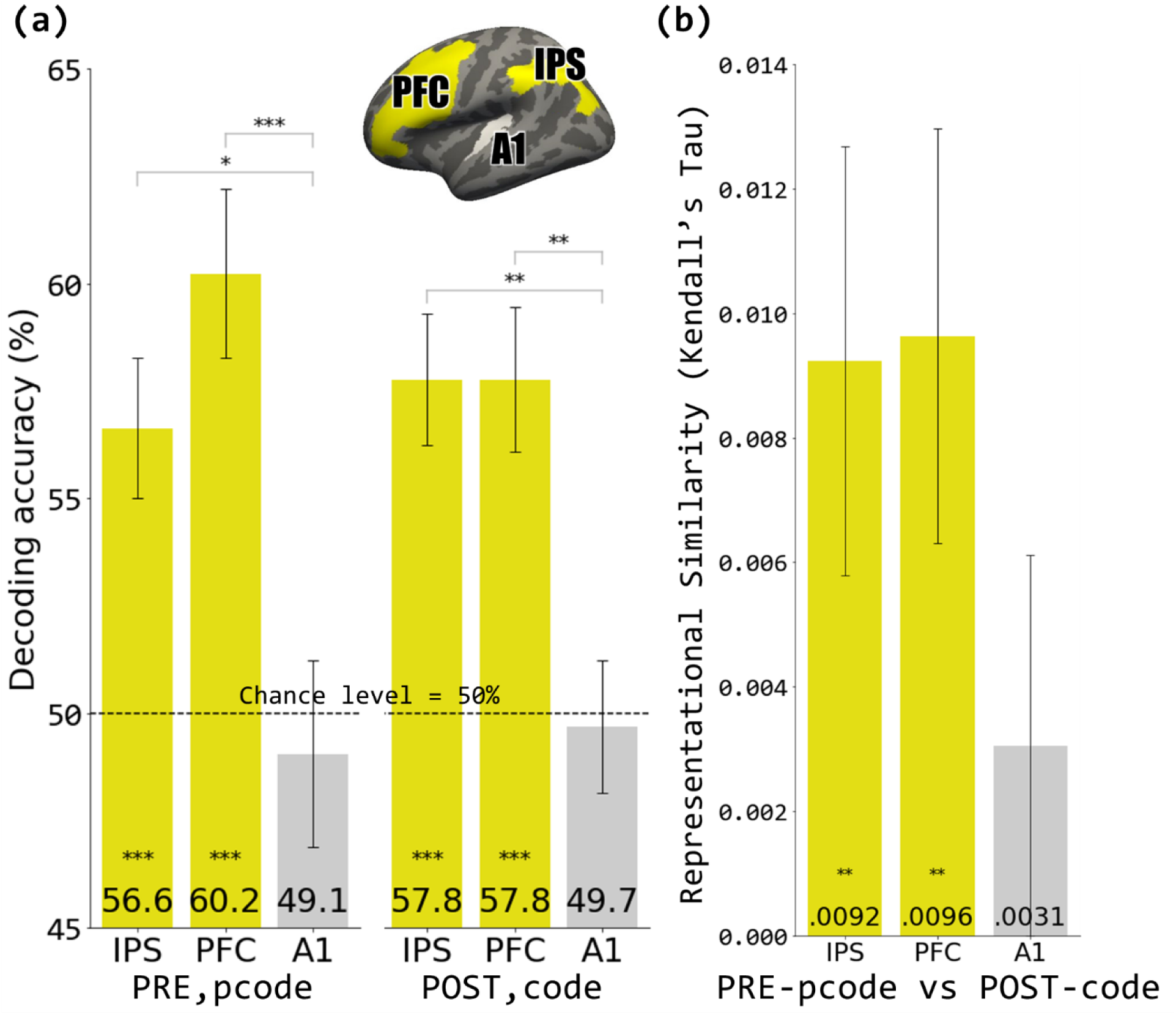
Multi-variate pattern analyses (MVPA). (a) binary decoding between FOR loop and IF conditional algorithms, with chance level of 50%. Inset: masks for the ROI search spaces used in MVPA. Within each ROI search space, participant-specific functional ROIs (fROI) were selected based on their respective code > memory control contrast in the POST scan. (b) Trial-wise representational similarity (Kendall’s Tau) between pseudocode (PRE scan) and code (POST scan) spatial responses. *p<0.05, **p<0.01, ***p<0.001.

Moreover, neural populations that respond to code after programming instruction already represented the “for”-vs-“if” distinction even prior to instruction. Parietal (IPS) and prefrontal (PFC) vertices that responded to code (over the memory control task) in the POST scan showed multivariate sensitivity to the distinction between “for” and “if” algorithms in the plain English pseudocode during the PRE scan (IPS W=190.0, p<0.001; PFC W=239.0, p<0.001; control region A1 W=97.0, p=0.62. Comparison against A1 IPS vs A1: W=180.5, p<0.05. PFC vs A1: W=223.5, p<0.001. Figure 3a). For-vs-if pseudocode can also be decoded during the POST scan (Supplementary Figure 4).

### Code “semantic” information in fronto-parietal reasoning network before and after programming instruction: representational similarity analysis (RSA)

Each participant saw the same algorithms presented as pseudocode during the PRE scan and as code during the POST scan. This allowed us to use RSA to probe the nature of algorithm representations in the IPS and the PFC and compare their representational content before and after programming instruction.

First, we asked whether vertices that go on to become involved in code comprehension represent similar information before and after instruction. Focusing on vertices that would eventually respond to programming code after instruction, we computed the representational similarity matrices (RSM) of neural responses to code algorithms (after instruction) and the similarity among neural responses to pseudocode algorithms (before instruction) in the IPS, the PFC, and the A1, which serves as a control region. The code RSM and the pseudocode RSM were significantly correlated in the IPS and the PFC, but not in the A1, suggesting shared information in the fronto-parietal network before and after instruction (Kendall’s Tau. IPS: mean=0.0092, SD=0.015, non-parametric permutation test p<0.001. PFC: mean=0.0096, SD=0.015, p<0.001. A1: mean=0.003, SD=0.014, p=0.16. Non-parametric sign tests IPS vs A1: p=0.10. PFC vs A1: p=0.064. Figure 3b).

Whole cortex search-light RSA revealed significant representational similarity between PRE-pseudocode and POST-code in the left IPS and PFC. Significant clusters were also observed in the left frontal pole and posterior temporal cortex, the latter falling within the language network (Supplementary Figure 5). These results suggest that fronto-parietal networks that represent programming algorithms after coding instruction already contain algorithm representations even before people learn to code. A small cluster was also observed in the primary visual cortex, suggesting presence of perceptual similarity among code and pseudocode.

We next used RSA to further probe the representational content in code-responsive fronto-parietal networks before and after instruction in more detail than the for/if distinction. We also used RSA to separate neural sensitivity to the semantics of code/pseudocode algorithms from the potentially confounded visual similarity of the code and pseudocode snippets.

We computed rich semantic similarity of the code and pseudocode snippets in terms of properties defined a priori to create the stimuli, as well as vector representations of the stimuli in large language models pre-trained on not only English texts but programming scripts. We also computed low-level perceptual similarity of the stimuli, and used principle component analysis (PCA) reduce these semantic and perceptual dimensions to a “meanings” feature and an “appearance” feature, respectively. We describe the details below.

The algorithms underlying Python code and pseudocode snippets varied systematically along five a priori defined dimensions. In addition to the distinction between for/if control structures, the other four dimensions include: the expression of the control structure (i.e., the iterable object for a “for” loop like the input string itself, or the unique characters in the input string; or the predicate of the “if” conditional, like the identity of a specific character in the input string, or the length of the input string), the data type of the “result” variable (character string or list), the operations performed on the “input” variable (e.g., reverse the input character string, repeat its first character three times, append two asterisks to its end), and what object resulted from the operation (e.g., a subset of the input, a single character, a list). We used these dimensions and the for/if distinction to compute the similarity of the code and pseudocode snippets to each other (Figure 4. Also see Supplementary Figure 6 & 7). We additionally derived “semantic” vector representations of code and pseudocode from two pre-trained large language models. We used CodeBERT to generate semantic vector representations of Python code stimuli (Z. Feng et al., 2020). CodeBERT has been trained on programming scripts in different programming languages and their documentations in GitHub repositories, and it has previously been shown to achieve high performance on code documentation generation and natural language code search. For pseudocode stimuli, we generated semantic vectors using OpenAI text-embedding-ada-002. The similarity among these vector representations was included as one of the features capturing the “semantic” information in the stimuli. Together, these resulted in six semantic features of code similarity. On the other hand, to capture the surface “appearance” of the code and pseudocode snippets, we counted the number of characters and the number of “words (tokens)” in each snippet.

**Figure 4.**
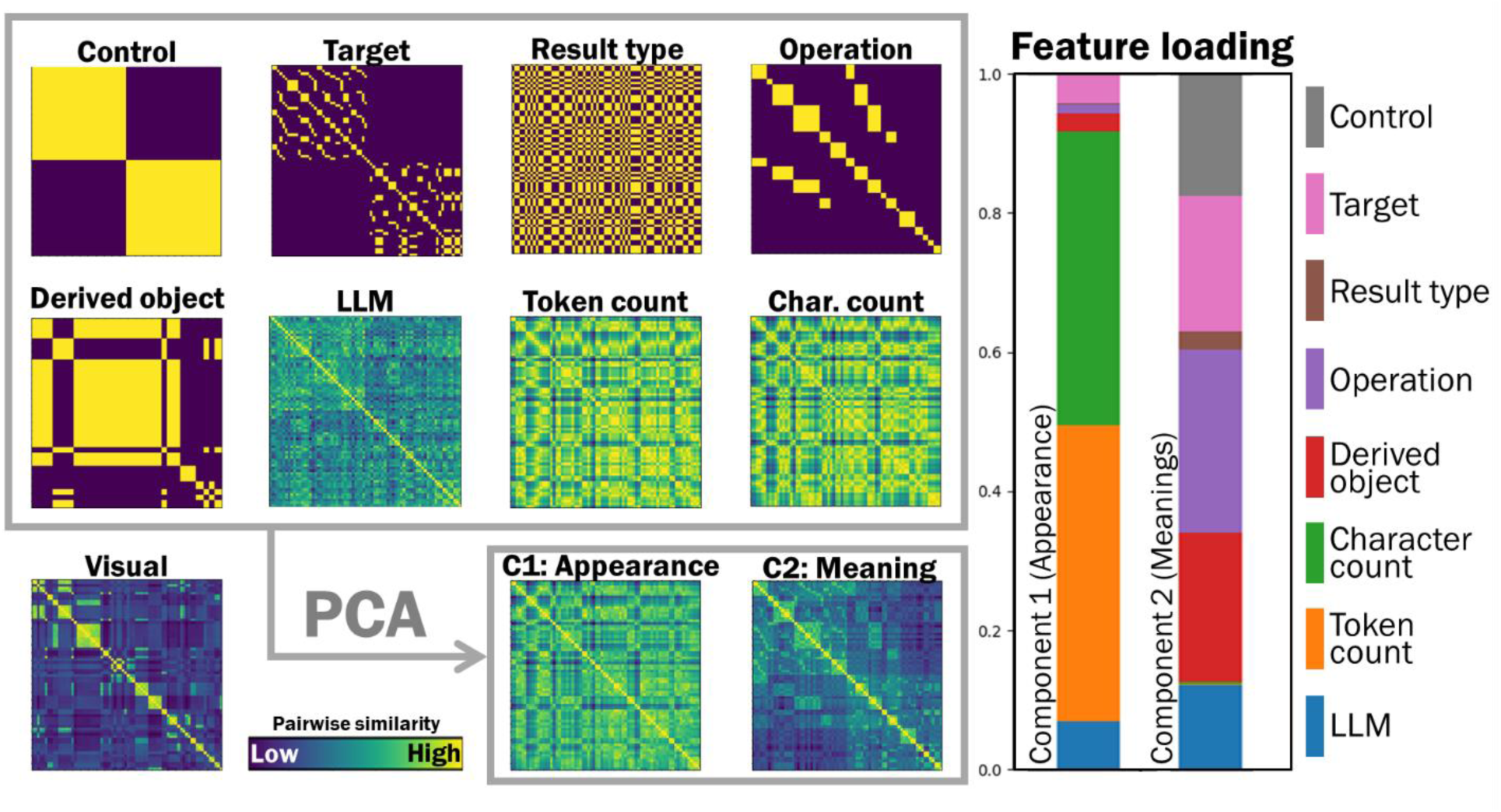
Example model matrices for representational similarity analysis (RSA). Left: the 9 representational similarity matrices (RSM) for the 9 features of the code stimuli. The “visual” RSM was left out as a confound for partial correlation, while the other 8 RSMs were submitted to principal component analysis (PCA) to derive 2 composite feature RSMs. Right: feature loadings for the two principal components. Component 1 is dominated by features related to the superficial appearance of the stimuli; Component 2 has contributions from features determining the “meaning” of the algorithms. This experiment included two batches of algorithms, each involved pseudocode and code versions. This figure shows the batch 1 code as an example. Please see Supplementary Figures 6 and 7 for the RSMs and PCA results for both batches and stimuli types.

In total, we derived eight feature dimensions, six of which were semantic- and two of which were appearance-related. From these eight features, we used PCA to identify two orthogonal components, where Component 1 captured the “appearance” of the stimuli, whereas Component 2 captured the “meanings” of the code algorithms (Figure 4, also see Methods). Finally, we calculated the perceptual visual similarity of the stimuli using pixel overlap. In the RSA, we used partial correlation to remove the effect of this “visual” covariate of no interest.

We performed a searchlight RSA analysis on the cortical surface, computing a partial correlation between the neural RSM and either the appearance component (Component 1) or meanings component (Component 2), with the visual pixel-overlap RSM included as a confound.

The neural representations of the “meanings” component were strongest in left-lateralized fronto-parietal reasoning network and fronto-temporal networks extending into inferior temporal and lateral occipital cortices (Figure 5, also see Supplementary Figure 8 for the results for POST pseudocode). By contrast, the neural representations of the “appearance” component were concentrated in the primary visual cortex. We then subtracted the partial correlation maps from the zero-order (no partial) correlation maps, which included pixel-overlap information. Through the subtraction, the primary visual cortex emerged as the sole area where the application of partial correlation substantially diminished the zero-order correlation between the neural RSMs and the feature RSMs (Supplementary Figure 9). These results suggest that the “meanings” of algorithms are represented in fronto-parietal and fronto-temporal network, and such “meanings” representation is beyond perceptual (visual) information. These representations show similar neural distributions before and after individuals learned how to program.

**Figure 5.**
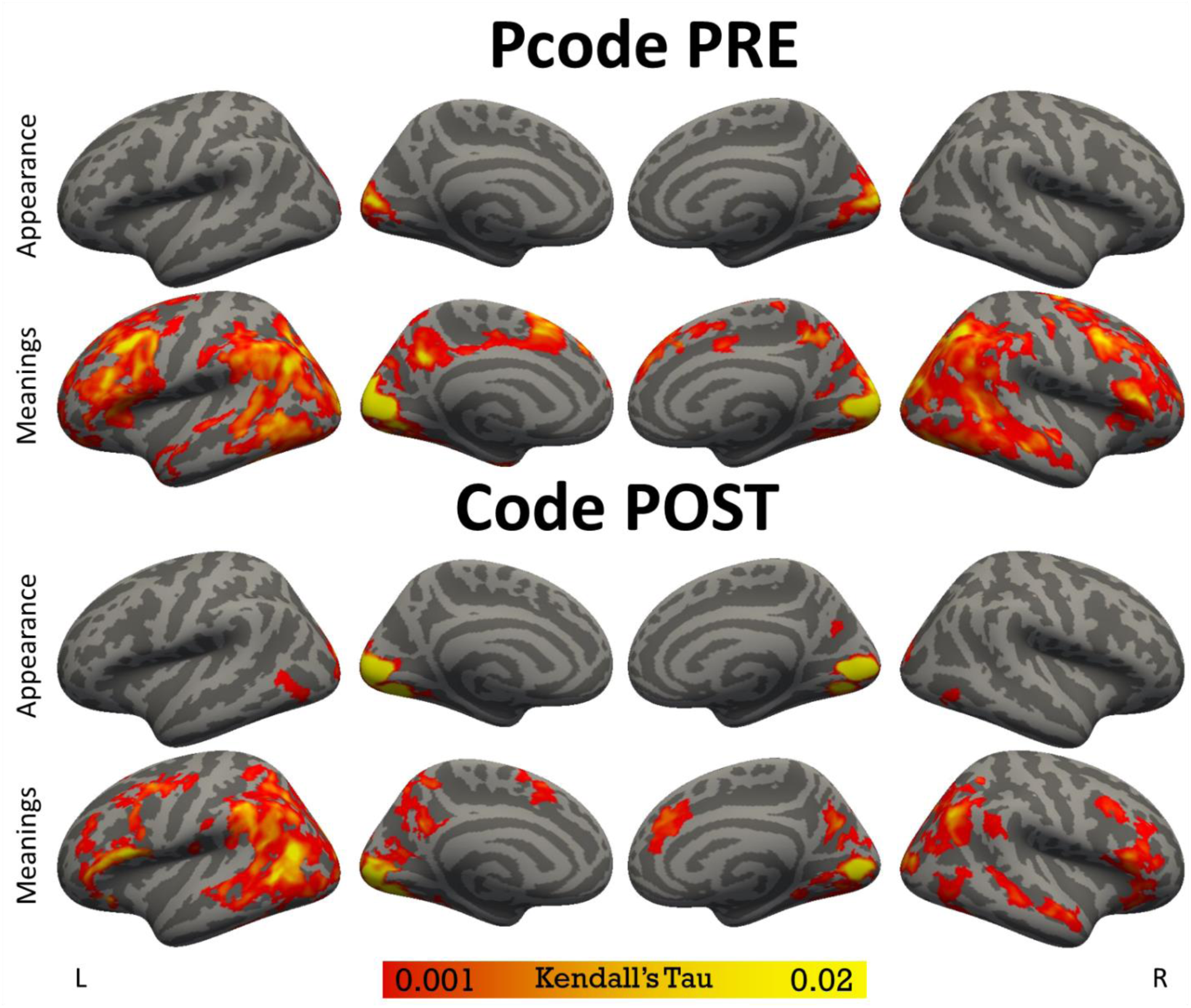
Whole-cortex searchlight representational similarity (Kendall’s Tau) between PRE pseudocode (top two rows) or POST code (bottom two rows) spatial activation patterns and either composite feature. Partial correlation was performed with visual RSM as the confound. Corrected for FWER with cluster-forming threshold of p<0.01 and cluster-wise p<0.05. For zero-order (no partial) correlation maps and the difference between zero-order and partial correlation maps, please see Supplement Figures 8 and 9.

## Discussions

We find that acquisition of programming skills involves rapid recycling or “reuse” of pre-existing representations of logical algorithms in a left-lateralized fronto-parietal network that supports logical reasoning, prior to code exposure. Previous studies found that expert programmers recruit a fronto-parietal logical reasoning network during code comprehension and activity patterns in this network are sensitive to a distinction between logical algorithms relevant to most programming languages, namely the “for” loops and “if” conditionals (Liu et al., 2020; Srikant et al., 2022). Here, we show that a similar network is recruited in beginning learners after their first Python programming course, after just 3.5 months of instruction.

Moreover, we find that code-relevant logical algorithms are represented in this fronto-parietal network even before programming instruction. Activity patterns in fronto-parietal neural networks that go on to respond to Python code after instruction distinguish between for vs. if algorithms in the same participants before instruction. In other words, we could decode the difference between plain English descriptions of executable “if” and “for” Python algorithms in a left-lateralized fronto-parietal reasoning network.

The present results support the idea that programming “recycles” pre-existing neural representations of logical algorithms (Dehaene, Al Roumi, Lakretz, Planton, & Sablé-Meyer, 2022; Liu et al., 2020). Our results are consistent with individual difference behavioral studies, which show that across programming languages, reasoning abilities – including fluid intelligence, working memory, mathematical problem-solving, logical deduction – are the strongest predictors of programming learning outcome in first-time programming students (Anderson et al., 1984; Farghaly & El-Kafrawy, 2021; Graafsma et al., 2023; McCoy & Burton, 1988; Prat et al., 2020; Shute, 1991). Reasoning abilities outperform linguistic abilities as predictors of programming learning outcomes (Farghaly & El-Kafrawy, 2021; Graafsma et al., 2023). Neuroimaging studies also find that reasoning tasks, such as logical deduction (e.g., “If X then Y. Not Y. Therefore, not X”), engage a specific fronto-parietal network (Coetzee & Monti, 2018; Holyoak & Lu, 2021; Monti & Osherson, 2012; Monti, Osherson, Martinez, & Parsons, 2007; Monti et al., 2009; Pischedda, Görgen, Haynes, & Reverberi, 2017; Wertheim & Ragni, 2020). Together with this prior evidence, the current results suggest that there is a specific relationship between fronto-parietal reasoning systems, and that code i.e., logical algorithm representations are recycled by programming instruction.

A key outstanding question concerns the ontogenetic origins of fronto-parietal algorithm representations. We measured sensitivity to these algorithms in young adults, who might have acquired them during their lifetime, prior to coding instruction, perhaps through formal education, as fronto-parietal reasoning systems are known for their protracted development and plasticity (Gogtay et al., 2004; Huttenlocher & Dabholkar, 1997). Alternatively, such representations might be innate or emerge early in life. Logical reasoning abilities are a human universal present across cultures in a wide variety of contexts, from programming to hunting and invention of new tools (Liebenberg, 2013; Pinker, 2022). Some recent evidence suggest that precursors of reasoning abilities emerge in early infancy. Preverbal infants are capable of disjunctive logical deduction (Nicolò Cesana-Arlotti, Kovács, & Téglás, 2020; Nicoló Cesana-Arlotti et al., 2018; Feiman, Mody, & Carey, 2022). There is also evidence fronto-parietal system are “online” from early infancy (Raz & Saxe, 2020) and engaged during simple rule learning in 8-month olds (Werchan, Collins, Frank, & Amso, 2016). Future neuroimaging work with children is needed to test the developmental origins of logical representations that go on to enable coding.

A further key direction for future work is to understand the neural changes that take place during programming instruction (Hishikawa et al., 2023; Hongo et al., 2022; Parnin, Siegmund, & Peitek, 2017). What distinguishes an expert coding brain from one that does not yet know how to code? One possibility is that neural representations of algorithms in fronto-parietal systems become elaborated or refined during programming instruction. Some evidence for this possibility comes from a recent study with coding experts (Ikutani et al., 2021). This study found better decoding of code algorithms in fronto-parietal systems of the participants with better performance on a programming task (Ikutani et al., 2021). Learning to code also involves forming a link between a novel cultural symbol system (e.g., Python syntax) and existing logical algorithm representations. Some evidence suggests that language systems may be involved in representing the surface form of code i.e., the language-like symbols (Coetzee & Monti, 2018; Liu, Wilson, & Bedny, 2024; Srikant et al., 2022). Interestingly, responses to Python code in fronto-parietal circuits are highly left lateralized, both in new learners and in expert programmers (Castelhano et al., 2018; Floyd et al., 2017; Ikutani et al., 2021; Ivanova et al., 2020; Krueger et al., 2020; Liu et al., 2020; Siegmund et al., 2014; Xu et al., 2021). A prediction to be tested in future work is whether connectivity between fronto-parietal and language systems are enhanced when people learn to code.

## Methods

### Participants

Participants were twenty-two Johns Hopkins undergraduates (11 women, 11 men, age range 18-24, mean age=19, SD=1.46) who were programming-naïve at the beginning of the study and enrolled in an introductory programming course “Gateway Computing: Python” at Johns Hopkins University. All participants completed the course in full. Participants took part in both MRI scanning sessions. None of the participants had any history of neurological conditions (screened through self-report). Informed consent was obtained from each participant in accordance with the Johns Hopkins Medicine Institutional Review Boards.

### Stimuli and task design

At the beginning of the semester, prior to the acquisition of programming knowledge, participants underwent an MRI scanning session (the “PRE” scan). At the end of the semester, after the last class of the programming course, another scanning session was administered (the “POST” scan).

During the PRE scan, participants completed a localizer task to identify their logical reasoning and language networks. During the same scan, participants also completed a “pseudocode” reading task and its corresponding control task. During the POST scan, participants completed another set of pseudocode reading task alongside a code reading task and their corresponding control tasks. The code reading task is conceptually identical to the task we administered in a previous study, with minor modifications (Liu et al., 2020). The “pseudocode” reading task was derived from the code reading task.

In the following paragraphs, we introduce the code and “pseudocode” stimuli, and their corresponding control stimuli (i.e., “scrambled code” or “nonsensical passage”).

#### Code stimuli: Python functions

A code comprehension trial consisted of three phases: “reading”, “input”, and “question”. During the “reading” phase, a Python function was presented. During the “input” phase, in addition to the function, an additional line was presented underneath the function to show the input to the function, based on which the participant derived the output. During the “question” phase, a single string of characters was presented, and participants answered whether it was the correct output of the function given the input. An example Python function is shown in Figure 1.

#### Code control stimuli: scrambled code

A code control stimulus contained all the words, symbols, and indentation structure of a Python function, but scrambled at the level of individual words and symbols such that it did not describe any executable algorithm. Participants were instructed to remember the lines in the scrambled function.

A code control trial also consisted of three phases. During the first “reading” phase, the scrambled function was presented. During the “input” phase, an additional scrambled line was presented underneath the scrambled function, which participants also had to remember. During the “question” phase, a single line of scrambled programming code elements was presented. Participant answered whether it was identical to one of the lines they just saw during the preceding phases. An example scrambled function is shown in Figure 1.

#### Pseudocode stimuli: plain English descriptions of algorithms

Each “pseudocode” stimulus was a short passage written in plain English to describe the algorithm implemented by a code stimulus (Python function). There is a one-to-one mapping between each element in a Python function and the words or phrases used in its corresponding pseudocode passage. An example pseudocode passage is shown in Figure 1.

The design and the task of a pseudocode reading trial was identical to a code reading trial. A pseudocode passage was presented during the “reading” phase. During the “input” phase, an additional sentence in the format “Now

<INPUT> is the word ‘_____’” was presented underneath the pseudocode passage, where “______” was replaced with a specific character string. Participants performed the algorithm described by the pseudocode based on the information contained in the additional sentence. During the “question” phase, a single character string was presented, and participants answered whether it was the correct result when the algorithm is performed. The pseudocode passages underwent multiple rounds of pilot-testing and modification to ensure comprehensibility and unambiguous wordings.

Please note that the “pseudocode” passages used in this experiment were different from the “pseudocode” used by software engineers. In the sense of software engineering, a pseudocode is a conceptual outline of a program. Although it does not have to be expressed in any particular programming language, usually it still resembles a programming script with the use of symbols, indentations, and non-linguistic expressions. In this experiment, we borrowed the term “pseudocode”, but not its engineering style, to refer to the plain English descriptions of algorithms.

#### Pseudocode control stimuli: nonsensical passages

Same as the code control task, the pseudocode control task was a memory task. However, instead of randomly scrambling all the words in a pseudocode passage (as have been done to create code control stimuli), we rearranged the words to create nonsensical passages containing grammatically correct sentences which, as a whole, did not implement an executable algorithm. Specifically, each nonsensical passage and the additional line presented during the “input” phase of the trial was created by pooling the words from two pseudocode passages (plus their additional lines) and manually creating sentences out of the pooled words. An example nonsensical passage is shown in Figure 1.

All the experiment stimuli were contained in an Open Science Foundation repository: (https://osf.io/2ncfm/), including Python functions, scrambled functions, pseudocode passages, and nonsensical passages, in both text format and picture format which was presented to participants inside the MRI scanner.

#### Localizer tasks and stimuli

The localizer task was identical to the one used in our previous study (Liu et al., 2020). It aimed to identify brain responses specific to formal logic, symbolic mathematics, and language comprehension, with each condition serving as a control for the others. This combined language/math/logic localizer task design was adapted from previous studies (Kanjlia et al., 2016; Monti et al., 2009, 2012).

During the language trials, participants determined whether two visually presented sentences, one in active voice and one in passive voice, conveyed the same meaning. For instance, they compared sentences like “*The child that the babysitter chased ate the apple*” with “*The apple was eaten by the babysitter that the child chased*”. During the math trials, participants assessed whether the variable X had an identical value across two equations. For example, they compared equations such as “*X minus twenty-five equals forty-one*” with “*X minus fifty-four equals twelve*”. For the formal logic trials, participants evaluated whether two logical statements were consistent, wherein both statements were valid inferences from each other and thus had the same truth table. For instance, they compared statements like “*If either not Z or not Y, then X*” with “*If not X, then both Z and Y*”.

Consistent with previous works (Kanjlia et al., 2016; Monti et al., 2009, 2012), in the current study, the language network was localized by contrasting the neural response during language trials against math trials. The logical reasoning network was localized by contrasting logic trials against language trials.

### Procedure

The PRE scan (pseudocode + localizer) and the POST scan (pseudocode + code) were separated for one semester (3.5 months). At least a day prior to both scanning sessions, participants took part in a behavioral practice session to familiarize with the stimuli.

Each of the code or pseudocode reading comprehension trial began with a 0.5-second fixation cross. The three phases of each trial – “reading”, “input”, and “question” – lasted for 20, 6, and 4 seconds, respectively, with a 0.5-second fixation cross between the “input” and the “question” phases. During the “question” phase, once the participant made a choice by pressing a button to indicate “true” or “false”, the remaining time of the question phase was skipped, and a 5-second inter-trial interval (ITI) began. Including the ITI, each trial took at most 36 seconds.

The PRE scan consisted of 9 runs. In each run, participants completed 8 pseudocode trials and 4 pseudocode control trials. The POST scan also consisted of 9 runs. In each run, participants completed 8 pseudocode trials, 8 code trials, 4 pseudocode control trials, and 4 code control trials. The pseudocode/code stimuli involved two conditions: half of them described “for” loop algorithms, where an operation is done repeatedly across a collection of items. The other half described “if” conditional algorithms, where an operation was done only if a certain criterion is met. Within each run, the correct answer to half of the trials was “yes”, while the correct answer for the other half was “no”. The order of trials was counterbalanced pseudo-randomly across participants.

During the PRE scan session, participants underwent 4 localizer runs, during which they completed the language/math/logic tasks. The beginning of each trial was marked by a 1-second fixation cross. For each pair of sentences/equations/logical statements, one of them appeared first, with the other following 3 seconds later. The whole pair remained visible on the screen for a duration of 16 seconds. Participants indicated their response as true or false by pressing either of two buttons. A 5-second ITI began after participants made a response.

Each of the 4 localizer runs comprised 8 trials of language, math, and logic tasks, respectively, and 6 randomly distributed 5-second rest periods. In half of the trials, the correct response was “yes”, and the order of trials was counterbalanced across participants.

### MRI data acquisition and preprocessing

All functional and structural MRI data were acquired at the F.M. Kirby Research Center of Functional Brain Imaging on a 3T Phillips dStream Achieva scanner. T1-weighted structural images were collected in 150 axial slices with 1 mm isotropic voxels using a magnetization-prepared rapid gradient-echo (MP-RAGE) sequence. Functional T2*-weighted BOLD scans were collected using a gradient echo planar imaging (EPI) sequence with the following parameters: 36 sequential ascending axial slices, repetition time (TR) = 2 seconds, echo time (TE) = 0.03 seconds, flip angle = 70°, field of view (FOV) matrix = 76 x 70, slice thickness = 2.5 mm, inter-slice gap = 0.5, slice-coverage FH = 107.5, voxel size = 2.53 x 2.47 x 2.50 mm, PE direction = L/R, first order shimming. Six dummy scans were collected at the beginning of each run but were not saved.

During functional scans, stimuli were presented with custom scripts written in PsychoPy (Peirce et al., 2019). The stimuli were presented visually on a rear projection screen, cut to fit the scanner bore. The participant viewed the screen via a front-silvered, 45°inclined mirror attached to the top of the head coil. The stimuli were projected with an Epson PowerLite 7350 projector. The resolution of the projected image was 1600 × 1200. Due to the hardware update in the scanning facility, for 4 out of the 22 participants, the visual stimuli were presented on an MRI-compatible display (Cambridge Research Systems BOLDscreen 32 UHD LCD displays) with a resolution of 3840 × 2160.

We used FSL, Freesurfer, the HCP workbench, and in-house Python and R scripts to analyze MRI data. During preprocessing, functional data were motion-corrected, high-pass filtered with a 128s cut-off, and re-sampled to the cortical surface for each participant using the standard Freesurfer pipeline (Dale, Fischl, & Sereno, 1999; Glasser et al., 2013; Smith et al., 2004). The surface data were then smoothed with a 6mm FWHM Gaussian kernel and prewhitened to remove temporal autocorrelation. Note that smoothing was performed on the surface, rather than in the volume, and that 6mm smoothing on the surface corresponds to approximate 3 mm smoothing in the volume (Hagler, Saygin, & Sereno, 2006). Cerebellar and subcortical structures were excluded from the analysis.

### Analysis

#### Univariate contrasts derived from general linear models (GLM)

In this analysis, we sought to reveal the neural networks with greater response to code or pseudocode than to their corresponding memory controls. For each participant, separate general linear models (GLMs) were constructed for the PRE pseudocode reading scan, the POST pseudocode+code reading scan, and the localizer scan. In each of these GLMs, we included a separate nuance regressor to model out time points with excessive motion, defined as time points in which frame displacement root mean squared (FDRMS) were greater than 1.5mm (Kim, Kanjlia, Merabet, & Bedny, 2017). White matter signal and cerebral spinal fluid (CSF) signal were also included in each of the individual subject GLMs as nuisance regressors.

In the GLM for the PRE pseudocode reading scan, 8 regressors, with temporal derivatives, were included after convolving with a canonical double-gamma hemodynamic response function. Three of the regressors modelled the 20-second stimulus “reading” phase of the three trial conditions (FOR pseudocode, IF pseudocode, nonsensical pseudocode memory control), respectively. Another 3 of the regressors modelled the “input” phase of the three conditions, respectively. One regressor modelled the “question” phase (proposed output and participant response), and the last regressor modelled the inter-trial interval (ITI).

In the GLM for the POST pseudocode+code reading scan, 14 regressors were included. 6 modelled the stimulus “reading” phase of each trial condition (FOR pseudocode, IF pseudocode, nonsensical pseudocode memory control, FOR code, IF code, scrambled code memory control), respectively. 6 modelled the input phase for each trial condition, 1 modelled the “question” phase, and 1 modelled the ITI.

In the GLM for the localizer scan, 4 regressors were included to model the language trials, logic trials, math trials, and the resting periods, respectively.

In all these GLMs, each run was modelled separately, and runs were combined within each participant using a fixed-effects model (Dale et al., 1999; Smith et al., 2004). Random-effects models were applied to conduct group-level analysis across participants. Models for group-level analyses were corrected for multiple comparisons using a nonparametric permutation test with a cluster-forming threshold of p<0.01, and family-wise error rate (FWER) controlled at p<0.05 (Eklund, Knutsson, & Nichols, 2019; Eklund, Nichols, & Knutsson, 2016; Nichols & Holmes, 2002; Winkler, Ridgway, Webster, Smith, & Nichols, 2014).

#### Multivariate pattern analysis (MVPA): “for” vs “if” decoding in regions of interest (ROIs)

In this analysis, we used support vector machine (SVM) classifiers to decode the neural representation of algorithms in three ROIs on the left hemisphere: the intra-parietal sulcus (IPS), the lateral prefrontal cortex (PFC), and the primary auditory cortex (A1) which served as a control.

The IPS and the PFC ROI search spaces were generated by combining the cortical parcels in the 400-parcel map reported by Schaefer et al. (2018) which encompassed the vertices activated in the group code reading > memory control contrast in our previous study (Liu et al., 2020). The A1 ROI was anatomically defined as the transverse temporal portion of a gyral-based atlas (Desikan et al., 2006; Morosan et al., 2001).

We used a functional ROI (fROI) approach for vertex selection within each ROI. The fROI approach addresses the variations in neural responses among individuals when performing a cognitive task (Kanwisher, McDermott, & Chun, 1997; Nieto-Castañón & Fedorenko, 2012). This approach allowed us to pinpoint the neural populations specific to each participant that were activated during the experimental conditions, all within a broader search space defined for the group as a whole. For each participant, in each ROI search space, the top 350 responsive vertices were selected based on the fixed-effect univariate code reading > memory control contrasts of the participant collected during the POST scan. The selected vertices constitute the fROI of that particular participant.

For each participant, for each stimulus set (PRE pseudocode, POST pseudocode, or POST code), a separate GLM was constructed to model the 20-second stimulus presentation phase of each trial. The resultant spatial patterns of z-statistics of the beta parameter estimation for each code/pseudocode reading trial was used to train a decoding classifier. Specifically, we trained and tested a support vector machine (SVM) classifier on the 350-vertex spatial patterns within each ROI search space for each participant. The SVM classifier was implemented by the PyMVPA toolbox, trained and tested using the default parameters (Hanke et al., 2009).

To ensure uniform signal strength throughout the MRI scanning sessions, data normalization was applied within each scanning run (Lee & Kable, 2018; Stehr, Garcia, Pyles, & Grossman, 2023). Within each run, in every fROI, the mean and standard deviation were calculated across trials and vertices and used for normalization, setting the mean to 0 and the standard deviation to 1. To avoid the dependency between trials from the same run artificially inflating the decoding accuracy, we performed a 9-fold leave-one-run-out cross-validation (Etzel, Valchev, & Keysers, 2011; Mumford, Davis, & Poldrack, 2014; Valente, Castellanos, Hausfeld, De Martino, & Formisano, 2021). In each cross-validation fold, the classifier was trained on the data from 8 out of the 9 task runs and tested on the left-out run. The resulting 9 accuracy values were averaged to derive the observed accuracy for a single participant in one fROI. Statistical significance of the group mean decoding accuracy was tested with Wilcoxon signed-rank test against chance level of 50%.

#### PRE-pseudocode vs POST-code representational similarity analysis (RSA)

The goal of this analysis was to explore whether there were consistent neural representations of the same algorithms before and after programming instructions.

In this study, participants saw the same 72 algorithms written as pseudocode during the PRE scan and as Python code during the POST scan. One participant was excluded from this analysis because he saw different pseudocode and code algorithms due to an error in experiment administration. For each of the remaining 21 participants, within each fROI, we created a symmetrical 72-by-72 representational similarity matrix (RSM) based on the spatial neural response patterns to PRE pseudocode and the POST code, respectively. The spatial neural response patterns were derived from the GLM created for the “for” vs “if” decoding, as introduced in the previous sub-section. Each entry in the RSM was the additive inverse of the Euclidean distance between the activation patterns for a pair of stimuli, such that a higher value indicated greater similarity between a pair of stimuli.

The correlation between the lower triangles of the PRE-pseudocode RSM and the POST-code RSM was measured with Kendall’s Tau (Kriegeskorte, Mur, Ruff, et al., 2008; Nili et al., 2014). Statistical significance was computed via a bootstrapping process where we computed the correlation between null RSMs with randomly permutated item labels. The permutation was performed 1,000 times to generate a distribution of null group mean correlation values across participants. The statistical significance of the real group mean was evaluated as the probability of observing a null value greater than the real value, under the null distribution.

We also conducted this PRE-pseudocode vs POST-code RSA across the whole cortical surface using the searchlight method (Kriegeskorte, Mur, & Bandettini, 2008; Su, Fonteneau, Marslen-Wilson, & Kriegeskorte, 2012). RSMs were constructed based on the spatial activation patterns in the “searchlight” surrounding each cortical vertex. A searchlight corresponding to a vertex encompassed all the vertices located within an 8mm diameter circle (as determined by geodesic distance) centered on that vertex (Glasser et al., 2013; Kriegeskorte, Goebel, & Bandettini, 2006). Searchlights containing any sub-cortical vertex were excluded.

To control for the family-wise error rate (FWER), we applied a cluster-based permutation correction method (Elli, Lane, & Bedny, 2019; Liu, Rapp, & Bedny, 2023; Musz, Loiotile, Chen, & Bedny, 2022; Regev, Honey, Simony, & Hasson, 2013; Schreiber & Krekelberg, 2013; Su et al., 2012). This correction involved a cluster-forming threshold of uncorrected p<0.01 and a cluster-wise FWER threshold of p<0.05 (Eklund et al., 2019; Eklund et al., 2016; Winkler et al., 2014).

#### Whole-brain searchlight RSA between neural activation and feature properties

In this analysis, we sought to identify the nature of the algorithm-related information encoded in the brain. We compared the second-order similarity structures (i.e., the representational similarity matrices, RSM) between neural responses and the stimuli along some feature dimensions. For each stimulus set (PRE pseudocode, POST pseudocode, POST code), a neural RSM was constructed in each searchlight on the cortical surface, as introduced in the previous sub-section. On the other hand, 9 RSMs were created to capture the similarity structure of the stimuli with regard to nine features: control structures (“for” or “if”), targets of the control structures (the loops in “for” functions can iterate through each letter in a character string, each item in a list, etc.; the conditionals in “if” functions can be based on the identity of the first letter in a string, the length of a string, etc.), data type of the result of the algorithm (list vs character string), operations taken within the control structures (e.g., repeat three times, add some characters, reverse), the objects derived from the operations (a single letter from the input string, a sub-string of the input string, etc.), the “semantics” of the algorithms represented in large language models (code: Microsoft CodeBERT; pseudocode: OpenAI text-embedding-ada-002), token count, character count, and the pixel layout of the visual presentation of the stimuli (Figure 4. See Supplementary Methods for more on the operational definitions of similarity along these dimensions).

Because any difference in any feature dimension led to a visual difference, the visual RSM was reserved as a confound for the correlation between neural RSMs and feature RSMs, which was addressed using partial correlation. The other eight feature RSMs were submitted to a principal component analysis (PCA) to derive two orthogonal composite RSMs through linear combination. Supplementary Table 2 showed the weights of each feature for each component. Consistent throughout the stimulus sets, component 1 was dominated by character count and token count, capturing the “appearance” of the stimuli, whereas component 2 involves high contributions from the features which defined the algorithms, thus capturing their “meanings”.

In each searchlight on the cortical surface, we first computed the zero-order (no partial) correlation between the neural RSM and either of the two component RSMs. Next, we computed the partial correlation with the visual RSM included as a confound (please refer to Supplementary Methods for implementation details). We also computed the difference between the zero-order and the partial correlation values to highlight the brain region(s) most affected by the exclusion of visual information. All the resulting brain maps were controlled for FWER using the same method and statistical threshold as the whole-cortex analysis described in the previous subsection.

## Supporting information

Supplemental results, methods, and figures

Table of activated brain regions

## Acknowledgements

We appreciate the support from the instructors of Gateway Computing: Python course – Dr. Siamak Ardekani, Dr. Kwame Kutten, Dr. Sara More, and Dr. Joanne Selinski. Their help in distributing participant recruitment notice was essential to this project. The design of our experiment stimuli also greatly benefited from the course material shared by the instructors. We also thank Catherine Chen, Carol Lu, Sangmita Singh, and Ziwen Wang for their contribution to behavioral data collection.

