## Supplemental results, methods, and figures for "Rapid “recycling” of logical algorithm representations in fronto-parietal reasoning systems following computer programming instructions"

### Supplementary Results

#### Behavioral results

In PRE session, a linear mixed-effect model was used to conduct post-hoc comparison between comprehension task and control task, with participants specified as the random effect. We found that participants had lower accuracy and longer response time for memory control task than for pseudocode comprehension task (control task accuracy mean=80.6%, SD=7.7%, compare against pcode comprehension $t=-2.09$, $p<0.05$; r.t. mean=2.1s, SD=0.23s, against pcode $t=10.81$, $p<0.001$). In POST scan, participants were also less accurate and slower on memory control (nonsensical passage) task than on pseudocode comprehension task (control task accuracy mean=81.1%, SD=12.2%, against pcode $t=-4.92$, $p<0.001$; r.t. mean=1.79s, SD=0.33s, against pcode $t=13.61$, $p<0.001$). As for code control task (scrambled), participants showed the same level of accuracy as code comprehension task, but slower in response (control task accuracy mean=86.6%, SD=8.2%, against code $t=1.45$, $p=0.15$; r.t. mean=1.48, SD=0.28, against code $t=2.24$, $p<0.05$).

#### For-vs-If decoding accuracy

| | IPS | PFC | A1 |
|---|---|---|---|
| **PRE pseudocode** | 56.6% (SD=7.5%) | 60.2% (SD=9.0%) | 49.1% (SD=9.9%) |
| **POST code** | 57.8% (SD=7.0%) | 57.8% (SD=7.7%) | 49.7% (SD=7.0%) |

#### MRI results for POST pseudocode

The network activated in the POST pseudocode > nonsensical passage memory control was significantly left-lateralized (network laterality index: mean=-0.35, SD=0.52, two-sided Wilcoxon signed-rank test against 0: $W=43$, $p<0.05$). It also overlapped more with the logic network than with the language network (dice coefficient. Overlap with logic: mean=0.27, SD=0.15, test against chance-level $p<0.001$. Overlap with language: mean=0.17, SD=0.11, test against chance-level $p<0.001$. Difference between logic and language assessed with a permutation test, $p<0.05$. Supplementary Figure 3).

The decoding accuracy between “for” and “if” pseudocode during the POST scan was significantly above chance in the PFC and the IPS, not in the A1 (IPS: mean=56.2%, SD=10.2%; $W=204$, $p<0.01$; PFC: mean=55.4%, SD=8.4%; $W=197.5$, $p<0.05$; A1: mean=48.4%, SD=6.8%, $p=0.86$). The decoding accuracy in the IPS and the PFC was both significantly greater than in the A1 (IPS vs A1: $W=189.0$, $p<0.01$. PFC vs A1: $W=176.0$, $p<0.01$. Supplementary Figure 4).

Whole-cortex searchlight representational similarity analysis (RSA) showed the same pattern as POST code – component 1 (the “appearance” component) was represented almost exclusively in the primary visual cortex, whereas the neural representations of component 2 (the “meanings” component) was wide-spread across the cortical surface, encompassing both the fronto-parietal reasoning network and the fronto-temporal language network, even after the application of partial correlation as an attempt to remove the effect of the visual RSM. The difference between zero-order and partial correlation revealed that the primary visual cortex is the region most significantly affected by partial correlation (Supplentary Figure 8 & 9).

### Supplementary Methods

#### Participants

Participants were informed to withdraw from the experiment should they drop the course. None of the 22 participants reported in this study withdrew from the experiment due to course dropping. Another individual participated the first MRI scan but not the second, his data were thus discarded.

#### Stimuli and task design

*Code stimuli: Python functions*

Each function contains five lines. The first line (`def fun(input):`) and the last line (`return result`) were the same across all functions. By systematically altering the elements in the remaining three lines, we created the collection of functions used this experiment. The second line initialized the variable "`result`", which was to be returned at the end of the function. The data type of "`result`" could be either a list or a character string. The third line specified the control structure, which could be either a "for" loop or an "if" conditional. For a "for" loop, examples for the item to be iterated through included the input character string of the function, a subset of the input, and a list generated by splitting the input string by a specific character (e.g., splitting the string `"2023-Dec-12"` by the character `"-"` yields the list `["2023", "Dec", "12"]`). For an "if" conditional, examples for the criteria for the conditional judgement included the length of the input, the identity of a particular character in the input, and the case of a particular character in the input. The fourth line was the body of the control structure which updated `result`. The new element added to `result` was derived by performing an operation on a component derived from `input`. The derived component could be the `input` string itself, a character in the string, a subset of the string, etc. The action could be repetition, case change, reversion of the order of the characters, etc. The additional line participants saw during the "input" phase was always in the format "`result = fun("……")`", where the example input "……" was replaced with a specific character string.

*Code control stimuli: scrambled code*

Same as real Python functions, each scrambled function consisted of exactly five lines. The first line (`input :def) fun()` and the last line (`result return`) were the same across all trials. The characters and symbols in the remaining three lines were pooled, scrambled, and redistributed. The resultant scrambled functions retain the indentation structure of, and hence are visually similar to, the real Python functions. Manual inspection was applied to ensure that none of the scrambled lines formed a legitimate Python expression.

*Pseudocode stimuli: plain English descriptions of algorithms*

There was a one-to-one mapping between each element in a Python function and the corresponding pseudocode passage. For example, in a "for" loop where some symbols were added to each character in the input string, the description was always in the format of "For EACH character in <input>, append N symbols to the character to make a new word" without any paraphrasing. The first line of pseudocode passages is always "<input> is a word" across the pseudocode reading trial. For all the pseudocode passages describing an "if" algorithm, the last line is always "Otherwise do nothing to <result>".

During the creation of pseudocode passages, some programming-specific terminologies were replaced with synonyms closer to everyday language, at the expense of the precision of the terminology. For example, "string" was replaced with "word", and "split the string by the token X" was replaced with "extract from the word the segments separated by the symbol X".

*Pseudocode control stimuli: nonsensical passages*

Each nonsensical passage and its corresponding additional line was created by pooling the words from two pseudocode passages (and their additional lines) and making sentences out of approximately half of the pooled words. Given the words in a single pseudocode passage, there was little flexibility in the structure of the grammatical sentences that could be created, especially given the constraint that the sentences should be incoherent from each other (i.e., cannot be understood as an executable algorithm). Due to the experiment design, participants saw twice as many pseudocode passages as control. Thus, creating one control stimulus out of two pseudocode passages yielded enough control stimuli while enhancing the variability of the sentences in nonsensical passages.

Similar to a pseudocode passage, all nonsensical passage begins with the sentence "<input> is in the character". In "if" nonsensical passages, the last sentence is always "Otherwise order the alphabetical list". Even though nonsensical passages do not describe "for" or "if" algorithms, "for" nonsensical passages still contain the word "for" but not the word "if", and vice versa for "if" nonsensical passages.

During the creation of nonsensical passages, care was taken to ensure that a nonsensical passage is visually similar to a pseudocode passage. Throughout the two MRI sessions, participants saw a total of 144 pseudocode passages (pcode) and 72 nonsensical passages (pctrl). Pseudocode and nonsensical passages were statistically identical in terms of the number of characters (pcode: mean=194.12, SD=24.24; pctrl: mean=196.86, SD=16.47; independent t-test: $t(214)=-0.86$, $p=0.39$).

*Grouping of participants and stimuli*

In this experiment, participants were divided into two groups, and the stimuli were also divided into two batches. The code and the pseudocode in the same batch describes the same algorithms. During the PRE scan (at the beginning of the semester), Group 1 participants saw Batch 1 pseudocode, and Group 2 participants saw Batch 2 pseudocode. During the POST scan (at the end of the semester), Group 1 participants saw Batch 2 pseudocode and Batch 1 code, whereas Group 2 participants saw Batch 1 pseudocode and Batch 2 code. Among the 22 participants, 9 were in Group 1, and 12 participants were in Group 2. Due to an error in experiment administration, one participant was in Group 1 during the PRE scan, but was in Group 2 during the POST scan. This participant was excluded from the representational similarity analysis (RSA) between PRE pseudocode and POST code.

#### Procedure

During the PRE scan, the trials were ordered such that none of the 3 conditions (FOR pseudocode, IF pseudocode, pseudocode control) appeared consecutively for more than 2 trials. During the POST scan, the trials were ordered such that none of the 6 conditions (FOR pseudocode, IF pseudocode, pseudocode control, FOR code, IF code, code control) appeared consecutively for more than 2 trials. To counter-balance the order of trial and task types seen by participants, upon participation, each participant was assigned 4 parameters to determine (1) the batch of stimuli they were going to see in each scanning session, (2) the order of trial conditions within the runs, (3) the assignment of the actual stimuli items to the trial conditions, and (4) which trials had “yes” (or “no”) as their correct answers.

#### A summary of the categories involved in this study

In this study, many categories have been introduced, and some of these categories partially nested within each other. Given the complexity of the experiment design, for the sake of clarification, in the following table, we listed all the categories pertinent to this study and the classes within each category.

*Supplementary Table 1*

<table>
<tr><th>CATEGORY</th><th>NUMBER</th><th>EXEMPLARS</th></tr>
<tr><td>Scan/session</td><td>2</td><td>PRE or POST</td></tr>
<tr><td>Stimuli type</td><td>2</td><td>Code or pseudocode. In this study, “pseudocode and/or code” is sometimes shortened as “pseudo/code”</td></tr>
<tr><td>Stimulus set</td><td>3</td><td>Combinations of scanning sessions and stimuli types: PRE pseudocode, POST pseudocode, POST code</td></tr>
<tr><td>Trial condition within each stimuli type</td><td>3</td><td>FOR, IF, memory control (FAKE)</td></tr>
<tr><td>Task type</td><td>4</td><td>pseudocode reading comprehension (FOR or IF conditions);<br>nonsensical memory control (FAKE condition);<br>code reading comprehension (FOR or IF conditions);<br>scrambled memory control (FAKE condition)</td></tr>
<tr><td>Phases in each trial</td><td>3</td><td>“reading (stimuli presentation)”, “input”, “question”</td></tr>
<tr><td>Batch of stimuli</td><td>2</td><td rowspan="2">Group 1 participants saw Batch 1 pseudocode in PRE session and Batch 2 pseudocode in POST session. For Group 2 participants, it was the opposite.</td></tr>
<tr><td>Group of participants</td><td>2</td></tr>
</table>

#### Analysis

##### *Cluster-based permutation correction for whole-cortex MVPA*

For multivariate pattern analysis (MVPA) – particularly representational similarity analysis (RSA) – conducted on the whole cortex using the searchlight method, we control for the family-wise error rate (FWER) using a cluster-based permutation correction method (Elli, Lane, & Bedny, 2019; Liu, Rapp, & Bedny, 2023; Musz, Loiotile, Chen, & Bedny, 2022; Regev, Honey, Simony, & Hasson, 2013; Schreiber & Krekelberg, 2013; Su, Fonteneau, Marslen-Wilson, & Kriegeskorte, 2012).

We conducted 100 label shuffles for the items (the 72 pseudocode or code stimuli). For each shuffle, one of the arguments for the correlation function was replaced with a null representational similarity matrix (RSM) generated based on the shuffled item labels. Correlation (Kendall's Tau) was carried out across the whole cortex to yield a brain map of null correlation values. Both the observed group-mean correlation map and each null map were subjected to a cluster-forming threshold of $p<0.01$ (based on one-sample t-tests of the mean correlation values across participants against 0). For every cluster, we calculated its strength-over-spread, which quantifies the average distance between each vertex within the cluster and the peak cluster, weighted by the decoding accuracy value of each vertex. We then recorded the maximum strength-over-spread value among the clusters in each null accuracy map, forming a null distribution comprising 100 maximum strength-over-spread values. A cluster in the observed map was considered significant if its strength-over-spread value exceeded the 95th percentile of the null distribution.

##### *Definitions of the representational similarity matrices (RSM) of stimuli features*

Five features were systematically manipulated to create the collection of algorithms used in this study: **control structures** ("for" or "if"), **targets** of the control structures (the loops in "for" functions can iterate through each letter in a character string, each item in a list, etc.; the conditionals in "if" functions can be based on the identity of the first letter in a string, the length of a string, etc.), **data type of the result** of the algorithm (list vs character string), **operations** taken within the control structures (e.g., repeat three times, add some characters, reverse), the **objects derived** from the operations (e.g., a single letter from the input string, a sub-string of the input string). The RSMs corresponding to these features are binary: if two stimuli share the same feature, the "similarity" between them is 1, otherwise 0.

Four continuous features of the stimuli and their RSMs were computed post-hoc: **semantic**, **character count**, **token count**, and **visual similarity**.

The **semantic similarity** between a pair of stimuli was defined as the **similarity between two stimuli in the representational space of large language models (LLM)**. Specifically, we relied on pre-trained large-language models. Either stimulus in the pair was tokenized and projected into the high-dimensional representational space of an LLM. The projection resulted in a vector representation of the stimulus, the similarity is thus computed as the cosine similarity between the vectors. For pseudocode passages, we used the "text-embedding-ada-002" engine trained by OpenAI (https://openai.com/blog/new-and-improved-embedding-model). For Python functions, we used Microsoft CodeBERT, a model trained on six programming languages including Python and specialized on the vector representation of programming scripts (https://github.com/microsoft/CodeBERT). Besides the vector representation, the LLM models also reported the number of tokens in a stimulus, which was used to construct the RSM for the token count feature.

The **character count** of a stimulus (which can be a Python function or a pseudocode passage) is the number of all characters in the stimulus. The **token count** of a stimulus is the number of all "words" (or any meaningful unit such as operators in Python like +=) in the stimulus. The similarity between two stimuli in terms of character count is defined as the smaller value divided by the larger value. For example, if one stimulus has 90 characters, and the other has 100, their similarity is 90/100 = 0.9. Similarity in terms of token count is defined in the same way.

To compute the **visual similarity** between two stimuli, we took the images presented to the participant, loaded them as single-channel (gray-scale), reshaped the two-dimensional images into one-dimensional vectors, then computed the Pearson correlation between the vectors. This feature captures how similar two stimuli are as they were seen by the participants during the scan (Fischer-Baum, Bruggemann, Gallego, Li, & Tamez, 2017).

As there were two batches of algorithms, there were two groups of the 5 RSMs corresponding to the 5 algorithm-defining features (control, target, result type, operation, derived object). Since there were code and pseudocode versions for each algorithm, there were four groups of the 4 RSMs computed post-hoc (semantic, character count, token count, visual). Please refer to Supplementary Figure 6 for all the involved RSMs.

##### *Using principle component analysis (PCA) to derive composite RSMs*

There is a certain degree of multicollinearity among these RSMs. For example, the correlation between the type of control structure and the target of the control is 0.3, the correlation between character count and token count can be as high as 0.8 for pseudocode. Therefore, rather than computing the correlations between these model RSMs with neural RSMs, we created composite RSMs by submitting these model RSMs to principal component analysis (PCA) followed by varimax rotation. We configured the PCA to yield two orthogonal components. As stated in the main text, the visual RSM was excluded from the PCA.

PCA was done for both batches of algorithms and both stimuli types (code or pseudocode) with each batch. Therefore, in total, four pairs of components were derived. Supplementary Table 2 showed the weights of each feature on each of the components. Supplementary Figure 7(b) illustrated the contribution of each feature to each of the components in terms of "loading", which is proportional to the square of the weights. Also, see Supplementary Figure 7(a) for all 8 component RSMs (2 batch x 2 stimuli type x 2 components).

*Supplementary Table 2*

| Type | Batch | Component | Control | Target | Result type | Operation | Derived object | Char count | Token count | LLM |
|---|---|---|---|---|---|---|---|---|---|---|
| Code | 1 | 1 | 0.012 | 0.228 | -0.053 | 0.156 | -0.095 | 0.666 | 0.669 | 0.141 |
| Code | 1 | 2 | 0.362 | 0.400 | 0.347 | 0.407 | 0.295 | 0.037 | 0.019 | 0.578 |
| Code | 2 | 1 | 0.000 | 0.157 | -0.012 | 0.119 | -0.099 | 0.690 | 0.680 | 0.115 |
| Code | 2 | 2 | 0.362 | 0.414 | 0.345 | 0.409 | 0.248 | 0.036 | 0.044 | 0.588 |
| Pseudocode | 1 | 1 | -0.032 | 0.134 | -0.063 | 0.187 | 0.082 | 0.685 | 0.680 | 0.067 |
| Pseudocode | 1 | 2 | 0.400 | 0.468 | 0.263 | 0.372 | 0.273 | 0.062 | 0.048 | 0.577 |
| Pseudocode | 2 | 1 | -0.033 | 0.115 | -0.016 | 0.260 | 0.313 | 0.628 | 0.613 | 0.224 |
| Pseudocode | 2 | 2 | 0.466 | 0.512 | 0.305 | 0.303 | 0.047 | 0.037 | 0.040 | 0.575 |

Please note that representational similarity analysis computes the correlation between pairs of RSMs, which is unaffected by linear translation or scaling of the RSMs. Thus, the exact values in RSMs are irrelevant; only the relative values matter. In an RSM, higher values indicate greater similarity between stimuli, with the critical information residing in the relative differences between entries.

#### *Partial correlation with visual RSM included as the confound*

Since any difference in the algorithms led to visual differences between a pair of stimuli, the component model RSMs inevitably represented the visual similarity structure to a certain extent. To explore whether regions with significant correlation values contain information that extends beyond visual similarity, we computed another RSA using partial correlation. We sought to mitigate the influence of the visual RSM from the correlation between each composite feature RSM and each neural RSM. Specifically, denoting X as the component RSM, Y as the neural RSM, and Z as the visual RSM, the zero-order (no partial) correlations between these variables were represented as $r_{XY}$, $r_{XZ}$, and $r_{YZ}$. Subsequently, the partial correlation between X and Y, while accounting for the presence of Z, was calculated as follows:

$$r_{XY.Z} = \frac{r_{XY} - r_{XZ} * r_{YZ}}{\sqrt{(1 - r_{XZ}{}^2) * (1 - r_{YZ}{}^2)}}$$

#### Supplementary figures

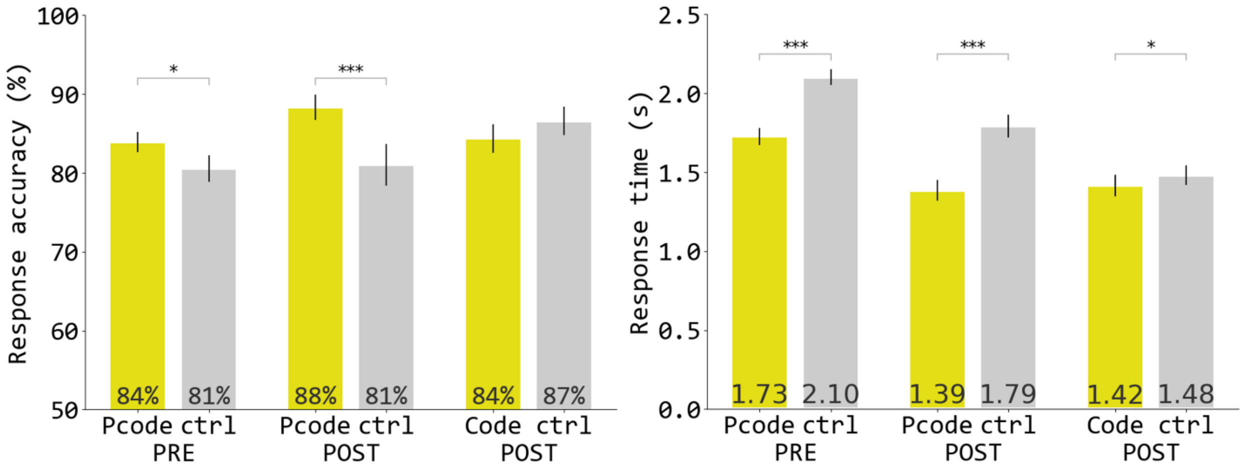


Supplementary figure 1: Behavioral performance of in-scanner tasks. "Pcode" means pseudocode. "Ctrl" means memory control task.

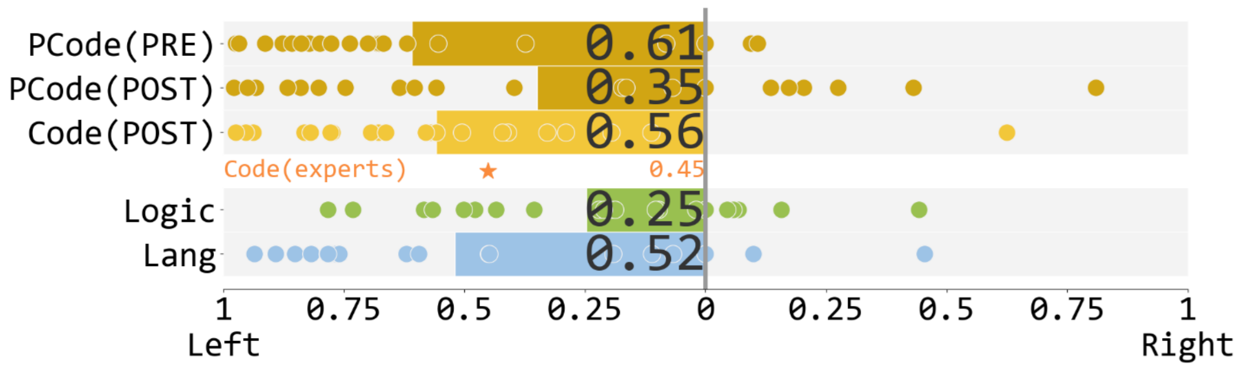


Supplementary Figure 2: Lateralization index of the univariate contrasts. The orange star denoted “Code (experts)” shows the lateralization index reported in our previous study (Liu et al., 2020).

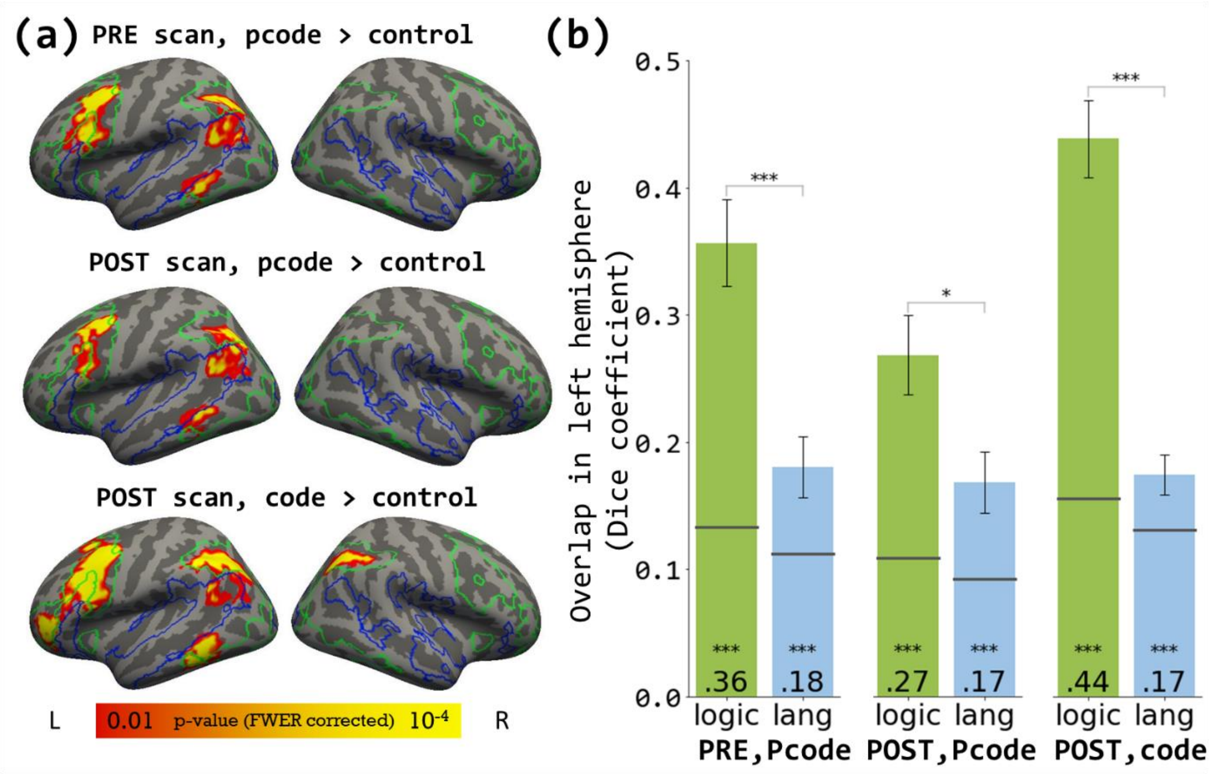


Supplementary Figure 3. Univariate responses. (a) From top row to bottom row: Pseudocode (pcode) > memory control contrast during the PRE scan; Pseudocode > memory control contrast during the POST scan; Code > memory control contrast during the POST scan. Logical reasoning network is outlined with green, language network in blue. All activation maps, including the localizers, were corrected for FWER with cluster-forming threshold of $p<0.01$ and cluster-wise $p<0.05$. (b) Overlap (Dice coefficient) between spatial activation patterns in (a) with logic or language (lang) networks localized within the same individuals. Error bars indicate standard error. Horizontal black lines on each bar indicate chance level overlap given the observed number of active vertices. Statistical significance of each bar was tested against respective chance level. ***$p<0.001$

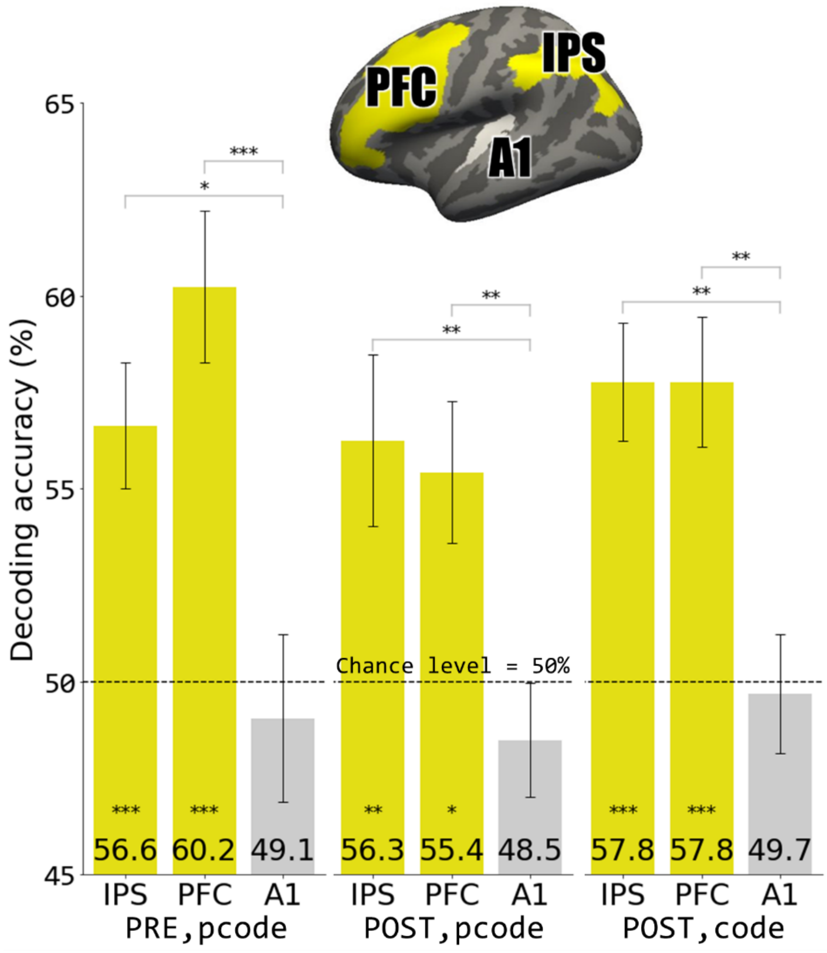


Supplementary Figure 4. Binary decoding between FOR loop and IF conditional algorithms, with chance level of 50%. Inset: masks for the ROI search spaces used in MVPA. Within each ROI search space, participant-specific functional ROIs (fROI) were selected based on their respective code > memory control contrast in the POST scan. *p<0.05, **p<0.01, ***p<0.001.

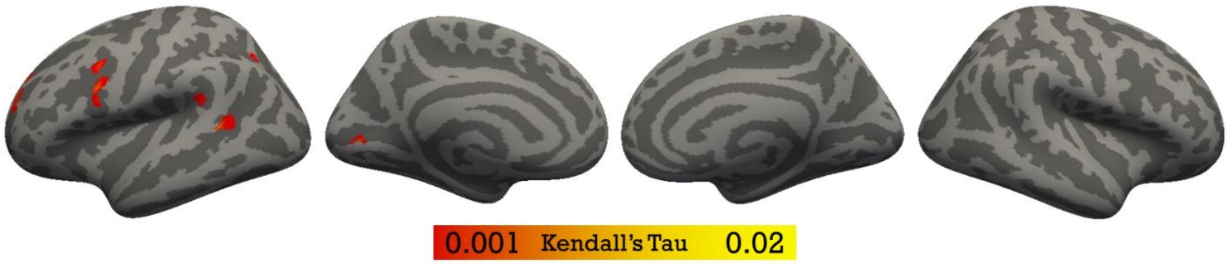


Supplementary Figure 5. Whole-cortex searchlight trial-wise PRE pseudocode vs POST code representational similarity. Corrected for FWER with cluster-forming threshold of p<0.01 and cluster-wise p<0.05.

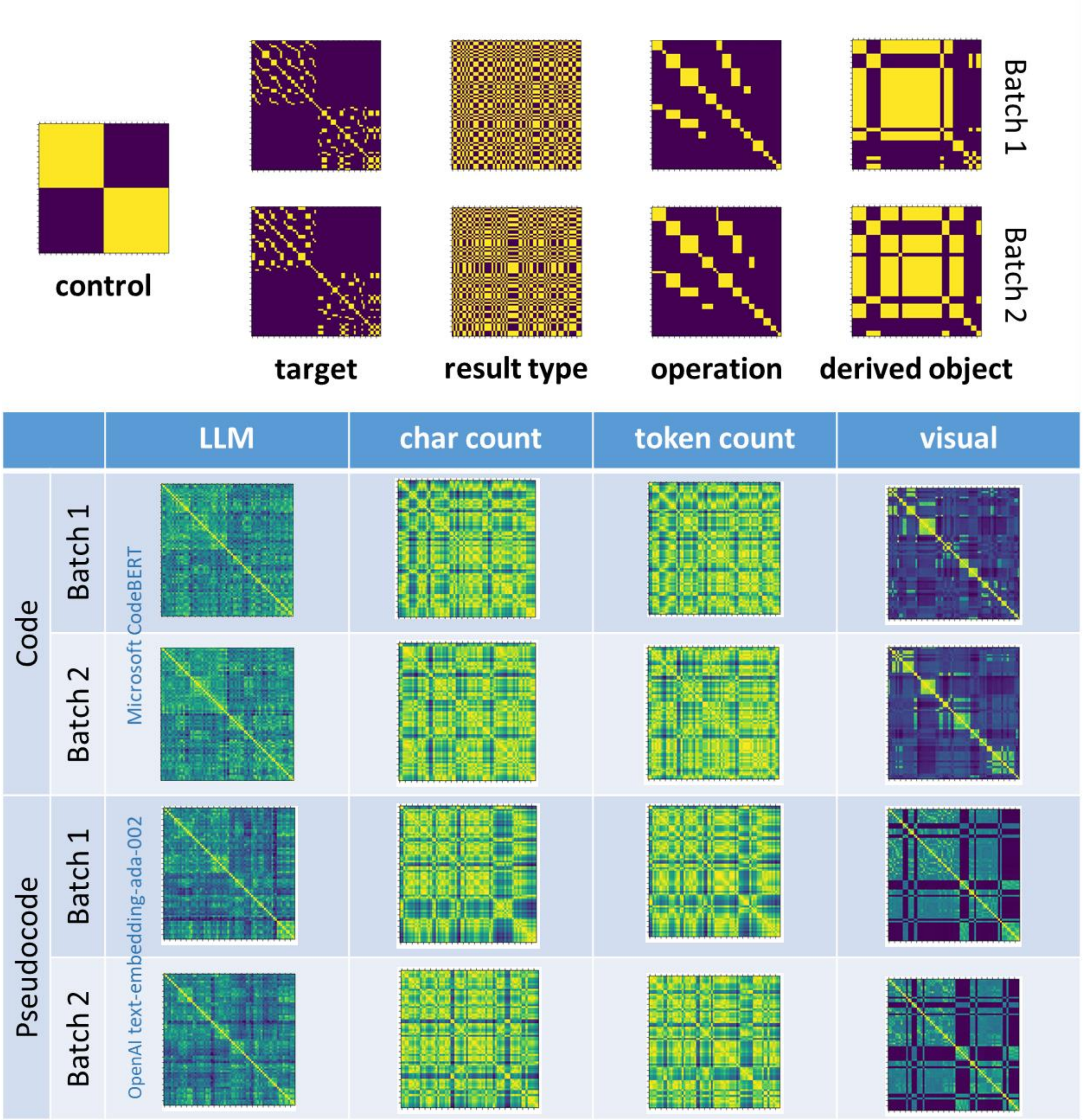


Supplementary Figure 6. Model matrices for representational similarity analysis (RSA). The categorical similarity matrices (control, target, result type, operation, derived object) are shared across code and pseudocode. The continuous similarity matrices (LLM, char count, token count, visual) were computed for each stimuli type (code or pseudocode) in each batch.

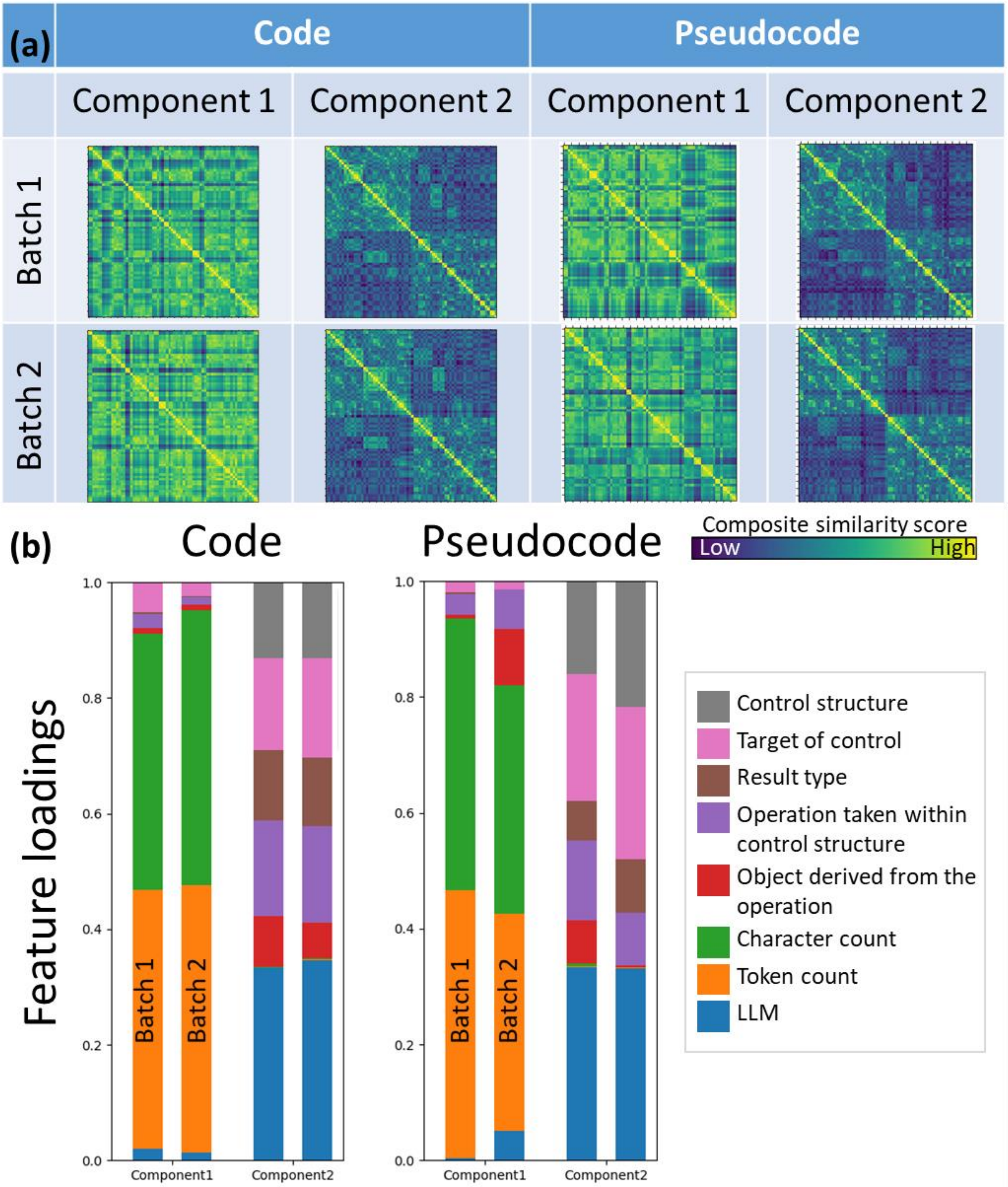


Supplementary Figure 7. (a) Composite feature matrices derived from PCA. (b) Feature loadings for the principal components.

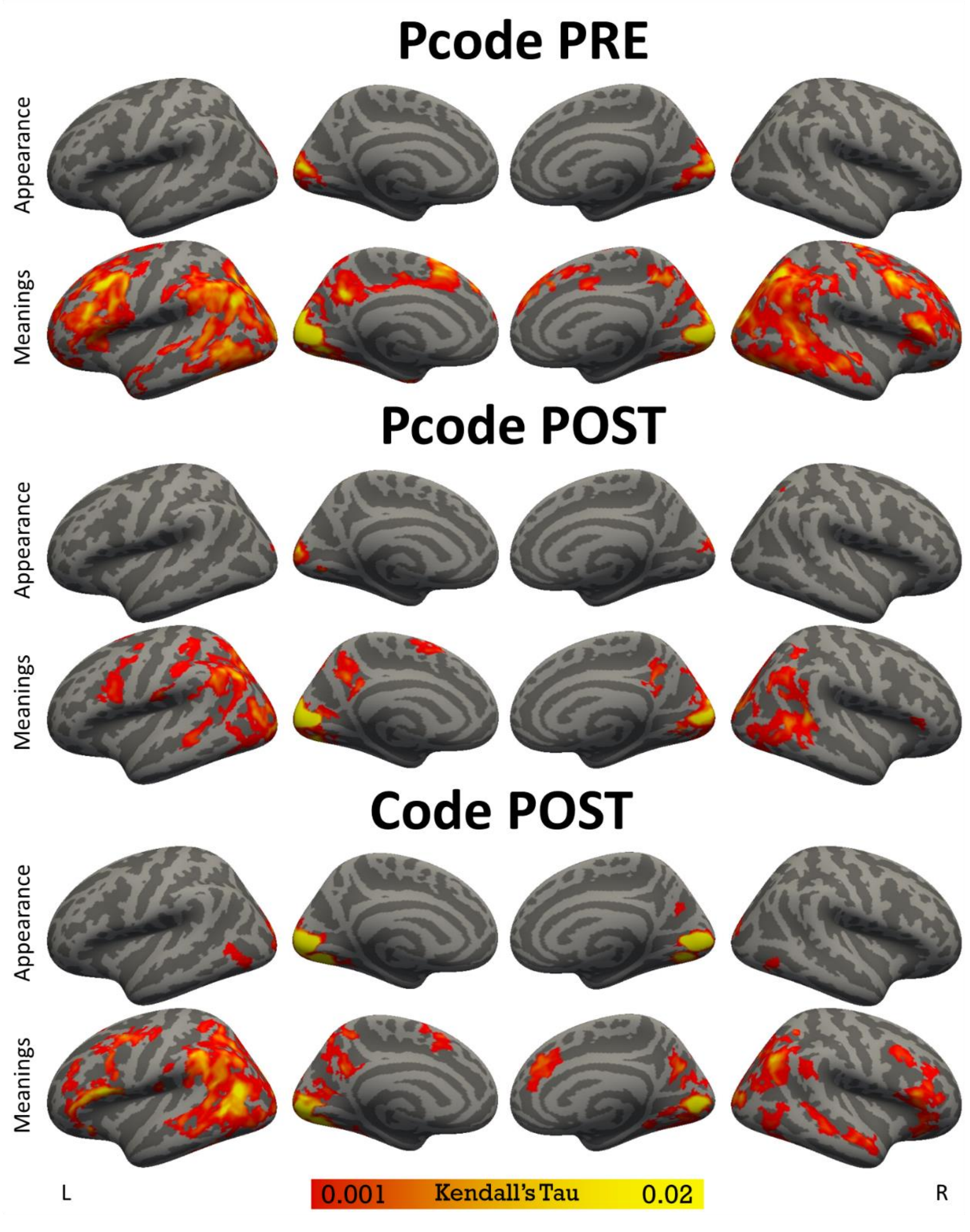


Supplementary Figure 8. Whole-cortex searchlight representational similarity (Kendall's Tau; zero-order correlation) between PRE pseudocode (top two rows), POST pseudocode (middle two rows), or POST code (bottom two rows) spatial activation patterns and either composite feature.

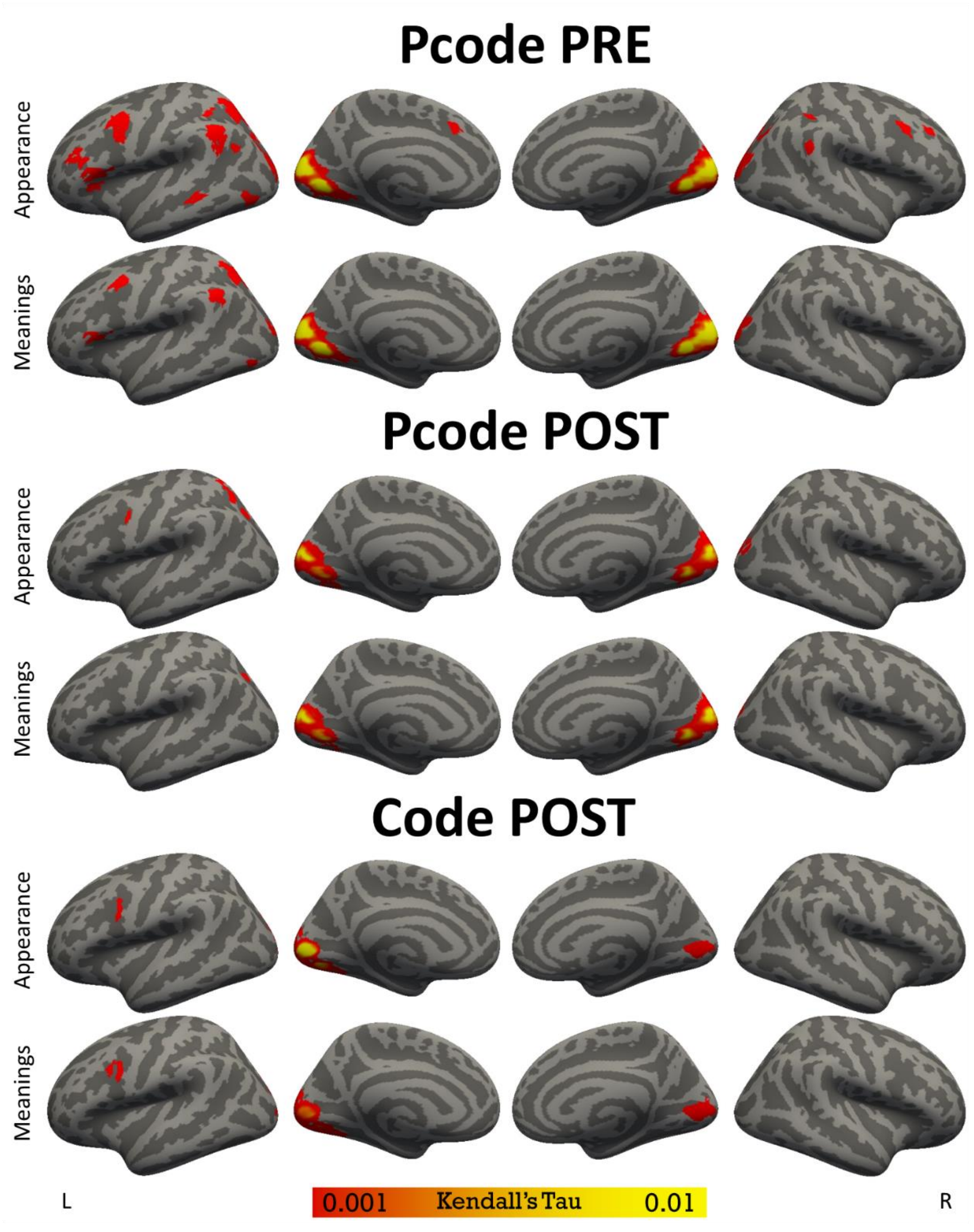


Supplementary Figure 9. Whole-cortex searchlight representational similarity (Kendall's Tau): difference between zero-order (Supplementary Figure 8) and partial correlations (Figure 5).

Elli, G. V., Lane, C., & Bedny, M. (2019). A Double Dissociation in Sensitivity to Verb and Noun Semantics Across Cortical Networks. *Cerebral Cortex*. doi:https://doi.org/10.1093/cercor/bhz014

Fischer-Baum, S., Bruggemann, D., Gallego, I. F., Li, D. S. P., & Tamez, E. R. (2017). Decoding levels of representation in reading: A representational similarity approach. *Cortex, 90*, 88-102. doi:https://doi.org/10.1016/j.cortex.2017.02.017

Liu, Y.-F., Rapp, B., & Bedny, M. (2023). Reading Braille by Touch Recruits Posterior Parietal Cortex. *Journal of Cognitive Neuroscience*, 1-24. doi:10.1162/jocn_a_02041

Musz, E., Loiotile, R., Chen, J., & Bedny, M. (2022). Naturalistic Audio-Movies reveal common spatial organization across "visual" cortices of different blind individuals. *Cerebral Cortex*, bhac048. doi:10.1093/cercor/bhac048

Regev, M., Honey, C., Simony, E., & Hasson, U. (2013). Selective and Invariant Neural Responses to Spoken and Written Narratives. *Journal of Neuroscience, 33*(40), 15978-15988. doi:10.1523/jneurosci.1580-13.2013

Schreiber, K., & Krekelberg, B. (2013). The Statistical Analysis of Multi-Voxel Patterns in Functional Imaging. *PLoS ONE, 8*(7), e69328. doi:https://doi.org/10.1371/journal.pone.0069328

Su, L., Fonteneau, E., Marslen-Wilson, W., & Kriegeskorte, N. (2012, 2-4 July 2012). *Spatiotemporal Searchlight Representational Similarity Analysis in EMEG Source Space.* Paper presented at the 2012 Second International Workshop on Pattern Recognition in NeuroImaging.
