## Supplementary material for "Rapid “recycling” of logical algorithm representations in fronto-parietal reasoning systems following computer programming instructions": Table of activated brain regions

Table 1. Clusters revealed in the group maps of univariate contrasts and multivariate pattern analyses. All clusters are corrected for FWER with cluster-forming threshold p<0.01 and cluster-wise threshold p<0.05.

| Cluster descriptions | peak MNI coordinates | | | Cluster size | | peak-p |
|---|---|---|---|---|---|---|
| | X | Y | Z | vertices | mm2 | |
| **Univariate code > control (POST)** | | | | | | |
| *Left hemisphere* | | | | | | |
| Intraparietal sulcus, angular gyrus | -35.2 | -66.7 | 45.1 | 1693 | 2578.04 | 1.24E-10 |
| Lateral prefrontal cortex: extending from precentral sulcus to frontal pole | -42.1 | 11.4 | 43.9 | 1629 | 3550.79 | 1.16E-10 |
| Precuneus | -4.5 | -71.5 | 40.6 | 1129 | 2012.91 | 2.16E-09 |
| Posterior middle temporal gyrus | -58.9 | -39.8 | -15.4 | 370 | 886.6 | 3.30E-07 |
| Calcarine sulcus | -4.9 | -78.3 | 4.7 | 326 | 807.95 | 1.45E-08 |
| Medial superior frontal gyrus | -6 | 35.9 | 44.8 | 151 | 439.86 | 1.65E-06 |
| *Right hemisphere* | | | | | | |
| Calcarine sulcus | 11.4 | -84.6 | 0.6 | 549 | 1589.92 | 3.23E-11 |
| Intraparietal sulcus, angular gyrus | 45.1 | -56.6 | 45.4 | 559 | 934.2 | 8.71E-07 |
| Precuneus | 3.9 | -65.4 | 38.9 | 379 | 692.25 | 6.73E-08 |
| Posterior cingulate gyrus | 4.4 | -30.7 | 29.3 | 257 | 397.79 | 4.74E-07 |
| Lingual gyrus | 13.2 | -78 | -10.8 | 151 | 407.32 | 8.12E-09 |
| **Univariate pcode > control (PRE)** | | | | | | |
| *Left hemisphere* | | | | | | |
| Intraparietal sulcus, angular gyrus, posterior superior temporal sulcus | -39.7 | -67.1 | 47.6 | 1255 | 1806 | 9.64E-09 |
| Precuneus | -5.4 | -59.3 | 36.6 | 769 | 1523.78 | 2.29E-07 |
| Lateral prefrontal cortex: middle frontal gyrus, inferior frontal sulcus | -40.8 | 21.4 | 41.1 | 590 | 1265.57 | 1.19E-06 |
| Posterior cingulate gyrus | -5.7 | -41.2 | 25.6 | 229 | 354.93 | 4.94E-06 |
| Posterior middle temporal gyrus | -63.8 | -42.7 | -3.3 | 219 | 501.36 | 6.80E-06 |
| *Right hemisphere* | | | | | | |
| Calcarine sulcus | 11.6 | -78.1 | 4.2 | 763 | 2048.15 | 1.78E-07 |
| **Univariate pcode > control (POST)** | | | | | | |
| *Left hemisphere* | | | | | | |
| Intraparietal sulcus, angular gyrus, posterior superior temporal sulcus | -39.7 | -67.1 | 47.6 | 1255 | 1806 | 9.64E-09 |
| Precuneus | -5.4 | -59.3 | 36.6 | 769 | 1523.78 | 2.29E-07 |
| Lateral prefrontal cortex: middle frontal gyrus, inferior frontal sulcus | -40.8 | 21.4 | 41.1 | 590 | 1265.57 | 1.19E-06 |
| Posterior cingulate gyrus | -5.7 | -41.2 | 25.6 | 229 | 354.93 | 4.94E-06 |
| Posterior middle temporal gyrus | -63.8 | -42.7 | -3.3 | 219 | 501.36 | 6.80E-06 |
| *Right hemisphere* | | | | | | |
| Calcarine sulcus | 11.6 | -78.1 | 4.2 | 763 | 2048.15 | 1.78E-07 |
| **RSA: PRE pcode vs POST code** | | | | | | |
| *Left hemisphere* | | | | | | |
| Posterior superior temporal sulcus | -47.5 | -49.9 | 7.4 | 72 | 83.59 | 0.00084 |
| Inferior precentral sulcus | -44.3 | 2.6 | 28.6 | 60 | 106.43 | 6.42E-05 |
| Anterior prefrontal cortex: middle frontal sulcus | -23.3 | 47.6 | 21.6 | 56 | 139.53 | 0.000102 |
| Inferior precentral sulcus | -36.3 | 6.3 | 34 | 38 | 52.9 | 0.000807 |
| Intraparietal sulcus | -27.9 | -64.3 | 41.2 | 38 | 35.26 | 0.000988 |
| Posterior lateral sulcus | -49.4 | -43.9 | 22.5 | 35 | 52.3 | 0.000907 |

| | | | | | | |
|---|---|---|---|---|---|---|
| Calcarine sulcus | -11.2 | -79.4 | 4.8 | 29 | 59.47 | 0.000571 |
| Anterior prefrontal cortex: superior frontal sulcus | -19 | 45.1 | 35.9 | 25 | 50.13 | 0.000575 |
| *Right hemisphere* | | | | | | |
| Dorsal paracentral gyrus | 15.7 | -41.1 | 75.1 | 47 | 96.01 | 0.000151 |
| **RSA: POST code "appearance" component (partial correlation with visual)** | | | | | | |
| *Left hemisphere* | | | | | | |
| Calcarine sulcus, lingual gyrus | -12.2 | -86.5 | 5.8 | 1426 | 3471.35 | 1.97E-09 |
| Posterior middle temporal gyrus | -54.2 | -61.4 | 2.7 | 151 | 233.69 | 0.000139 |
| *Right hemisphere* | | | | | | |
| Calcarine sulcus, lingual gyrus | 7.6 | -74.6 | 6 | 809 | 2202.37 | 4.27E-09 |
| Lateral inferior temporal gyrus | 49.1 | -60.9 | -2.5 | 65 | 117.77 | 0.000305 |
| Medial occipito-parietal sulcus | 16.7 | -63.8 | 31 | 57 | 108.46 | 8.31E-05 |
| **RSA: POST code "appearance" component (difference between zero-order and partial correlation)** | | | | | | |
| *Left hemisphere* | | | | | | |
| Calcarine sulcus, lingual gyrus | -9.5 | -91 | 6.1 | 1147 | 2826.23 | 4.46E-11 |
| Inferior precentral sulcus | -39 | -0.1 | 31.2 | 55 | 90.38 | 0.000946 |
| *Right hemisphere* | | | | | | |
| Calcarine sulcus, lingual gyrus | 5.3 | -75.5 | 5.6 | 177 | 526.68 | 2.01E-06 |
| **RSA: POST code "meanings" component (partial correlation with visual)** | | | | | | |
| *Left hemisphere* | | | | | | |
| The union of multiple sub-regions in posterior lateral occipito-temporo-parietal cortex and primary visual cortex | -15.6 | -84.5 | -12.5 | 6380 | 12034.69 | 9.29E-10 |
| Inferior frontal gyrus | -50.3 | 35.3 | -0.7 | 549 | 1305.6 | 2.03E-05 |
| Middle frontal gyrus | -40.3 | 5.9 | 52.4 | 594 | 1264.02 | 6.99E-05 |
| Medial superior frontal gyrus | -7.4 | 14.8 | 54.4 | 169 | 415.8 | 4.91E-06 |
| Anterior middle frontal gyrus | -38.9 | 39.5 | 29.2 | 164 | 376.79 | 7.91E-05 |
| Lateral occipito-temporal sulcus | -39.4 | -50.6 | -21.8 | 148 | 314.48 | 0.000138 |
| Subcentral cortex | -43.3 | -19.5 | 14.7 | 85 | 174.7 | 4.67E-05 |
| Superior frontal gyrus | -14.2 | 22 | 57.9 | 63 | 152.85 | 1.24E-06 |
| Anterior insula sulcus | -37.4 | 25.9 | -10.2 | 51 | 141.2 | 0.000273 |
| Supramarginal gyrus | -58.7 | -24.1 | 25.5 | 34 | 42.34 | 0.001548 |
| *Right hemisphere* | | | | | | |
| Intraparietal sulcus, angular gyrus, posterior superior temporal sulcus, superior occipital gyrus, middle occipital gyrus | 48.5 | -58.1 | 38.6 | 1658 | 3450.81 | 1.83E-07 |
| Inferior frontal cortex | 42.7 | 30.6 | -14.4 | 794 | 1757.77 | 2.49E-06 |
| Calcarine sulcus, lingual gyrus | 8.9 | -79.2 | 3.7 | 648 | 1829.18 | 3.47E-06 |
| Anterior cingulate cortex | 8.2 | 32.3 | 31.6 | 319 | 694.29 | 0.000172 |
| Lateral occipito-temporal cortex | 46.4 | -60.7 | -1.3 | 283 | 508.05 | 0.000159 |
| Lateral superior temporal gyrus | 53.1 | -1.7 | -17.2 | 229 | 525.94 | 2.02E-06 |
| Superior temporal sulcus | 49.7 | -31.3 | -7.8 | 209 | 392.28 | 2.84E-06 |
| Precuneus | 14 | -63.6 | 30.4 | 177 | 374.66 | 9.64E-06 |
| Inferior precentral sulcus | 36.7 | 7.7 | 36.8 | 160 | 288.46 | 9.87E-05 |
| Supramarginal gyrus | 59.7 | -42.1 | 26.7 | 154 | 238.03 | 0.000225 |
| Lingual sulcus | 26 | -41 | -10.8 | 89 | 210.6 | 6.04E-06 |
| Intraparietal sulcus | 34.2 | -49.4 | 51.8 | 50 | 71.46 | 0.000471 |

| | | | | | | |
|---|---|---|---|---|---|---|
| **RSA: POST code “meanings” component (difference between zero-order and partial correlation)** | | | | | | |
| *Left hemisphere* | | | | | | |
| Calcarine sulcus, lingual gyrus | -10.2 | -92.3 | 5.8 | 1200 | 2950.68 | 2.72E-10 |
| Inferior precentral sulcus | -39 | -0.1 | 31.2 | 96 | 155.35 | 0.000525 |
| *Right hemisphere* | | | | | | |
| Calcarine sulcus, lingual gyrus | 3.7 | -75.9 | 4.2 | 229 | 683.72 | 1.73E-07 |
| **RSA: PRE pcode “appearance” component (partial correlation with visual)** | | | | | | |
| *Left hemisphere* | | | | | | |
| Calcarine sulcus, lingual gyrus, cuneus gyrus | -3.2 | -89.1 | 8.1 | 687 | 1670.55 | 5.07E-08 |
| *Right hemisphere* | | | | | | |
| Calcarine sulcus, lingual gyrus, cuneus gyrus | 15.2 | -79.3 | 8.3 | 615 | 1720.59 | 1.26E-07 |
| **RSA: PRE pcode “appearance” component (difference between zero-order and partial correlation)** | | | | | | |
| *Left hemisphere* | | | | | | |
| Calcarine sulcus, lingual gyrus, cuneus gyrus | -11.3 | -98.8 | 12.1 | 1396 | 3582.47 | 1.79E-09 |
| Intraparietal sulcus | -25 | -63.7 | 48.7 | 343 | 582.39 | 2.31E-05 |
| Inferior precentral sulcus | -39.3 | 6.2 | 44.3 | 272 | 484.58 | 8.93E-05 |
| Supramarginal gyrus | -50.6 | -51.9 | 30.3 | 262 | 398.19 | 5.89E-05 |
| Inferior frontal gyrus | -53.8 | 25.4 | 11.9 | 173 | 368.61 | 5.38E-06 |
| Middle frontal gyrus | -42.8 | 34.4 | 23.8 | 107 | 204.32 | 0.001535 |
| Lateral inferior occipital cortex | -45 | -73.7 | -9.2 | 79 | 118.8 | 0.00032 |
| Superior temporal sulcus | -53.9 | -33.3 | -9.2 | 64 | 126.48 | 0.001594 |
| Anterior superior insula sulcus | -31.7 | 27.6 | 6.6 | 61 | 100.18 | 0.001035 |
| Intraparietal sulcus | -34.5 | -44.7 | 38 | 63 | 80.44 | 0.002227 |
| Posterior superior temporal sulcus | -39.2 | -58.7 | 25.2 | 43 | 48.62 | 0.001637 |
| Anterior cingulate cortex | -10.4 | 19.8 | 36.7 | 38 | 72.07 | 0.001269 |
| *Right hemisphere* | | | | | | |
| Calcarine sulcus, lingual gyrus, cuneus gyrus | 4.7 | -88.8 | 13.3 | 1281 | 3641.89 | 2.42E-10 |
| Angular gyrus | 33 | -71.7 | 30.4 | 71 | 167.76 | 0.000153 |
| Intraparietal sulcus | 40.4 | -42.2 | 35.8 | 72 | 65.21 | 0.000633 |
| Inferior frontal sulcus, inferior precentral sulcus | 36.8 | 13.4 | 35.2 | 62 | 109.1 | 0.00027 |
| Middle frontal cortex | 33.7 | 30 | 34.5 | 50 | 64.24 | 0.000301 |
| Supramarginal gyrus | 58.7 | -44.8 | 20.2 | 52 | 80.21 | 0.001219 |
| **RSA: PRE pcode “meanings” component (partial correlation with visual)** | | | | | | |
| *Left hemisphere* | | | | | | |
| The union of multiple sub-regions in posterior lateral occipito-temporal-parietal cortex and primary visual cortex | -9 | -90 | 6.4 | 6304 | 12493.87 | 7.83E-10 |
| The union of multiple sub-regions in lateral frontal cortex | -50 | 21.1 | 6.4 | 2610 | 5394.93 | 8.66E-07 |
| Dorsal medial superior frontal cortex, middle cingulate cortex, precuneus | -10.1 | 15.1 | 63.5 | 1457 | 2843.48 | 1.25E-06 |
| Anterior prefrontal cortex, orbitofrontal cortex | -23.4 | 54.2 | 24.7 | 601 | 1622.81 | 2.15E-06 |
| Temporal pole | -32.6 | -5 | -41.8 | 429 | 1309.61 | 2.65E-05 |
| Lateral superior temporal gyrus | -64.8 | -17.7 | -0.9 | 112 | 232.77 | 0.000155 |
| Precuneus | -8.6 | -62.9 | 57.8 | 104 | 154.83 | 0.000145 |

| | | | | | | |
|---|---|---|---|---|---|---|
| *Right hemisphere* | | | | | | |
| The union of multiple sub-regions in posterior lateral occipito-temporal-parietal cortex and primary visual cortex | 15 | -83.5 | 7.2 | 6528 | 13468.27 | 2.85E-08 |
| The union of multiple sub-regions in lateral frontal cortex | 40.8 | 6.7 | 48.9 | 2666 | 5826.65 | 3.72E-07 |
| Precuneus | 8.7 | -60.5 | 44 | 385 | 606.2 | 3.14E-05 |
| Central sulcus | 43.8 | -18.8 | 37.6 | 112 | 173.78 | 0.000271 |
| Medial superior frontal gyrus, middle/anterior cingulate cortex | 8.5 | 14.1 | 47.5 | 100 | 198.52 | 0.000234 |
| Anterior superior insula cortex | 32.8 | 19.7 | 4.7 | 105 | 163.1 | 0.000374 |
| Lateral superior temporal gyrus | 62.7 | -3.5 | -1.2 | 73 | 199.51 | 0.000179 |
| Middle/posterior cingulate cortex | 4 | -2.8 | 41.5 | 83 | 130.6 | 0.000539 |
| Orbitofrontal cortex | 34.9 | 36.7 | -7.8 | 63 | 170.98 | 0.000371 |
| Medial occipito-parietal sulcus | 17 | -57.4 | 12.1 | 53 | 103.24 | 0.000734 |
| Paracentral cortex | 3 | -36.7 | 63.7 | 47 | 112.8 | 0.000331 |
| **RSA: PRE pcode "meanings" component (difference between zero-order and partial correlation)** | | | | | | |
| *Left hemisphere* | | | | | | |
| Calcarine sulcus, lingual gyrus, cuneus gyrus | -7.1 | -98 | 5.8 | 1134 | 2967.73 | 2.20E-09 |
| Intraparietal sulcus | -25 | -63.7 | 48.7 | 274 | 460.12 | 3.54E-05 |
| Supramarginal gyrus, angular gyrus | -50.6 | -51.9 | 30.3 | 134 | 204.03 | 0.000251 |
| Inferior precentral sulcus, middle frontal gyrus | -38.5 | 5.5 | 42 | 113 | 196.76 | 0.000371 |
| Inferior frontal gyrus | -53.8 | 25.4 | 11.9 | 108 | 227.9 | 8.59E-05 |
| Inferior occipital cortex | -45 | -73.7 | -9.2 | 35 | 56.33 | 0.001422 |
| *Right hemisphere* | | | | | | |
| Calcarine sulcus, lingual gyrus, cuneus gyrus | 4.7 | -88.8 | 13.3 | 1111 | 3139.21 | 6.68E-10 |
| **RSA: POST pcode "appearance" component (partial correlation with visual)** | | | | | | |
| *Left hemisphere* | | | | | | |
| Calcarine sulcus, cuneus gyrus | -12.1 | -93.6 | 3.7 | 327 | 824.16 | 1.37E-05 |
| Lingual gyrus | -7.7 | -76.1 | -4.8 | 31 | 85.48 | 7.23E-05 |
| *Right hemisphere* | | | | | | |
| Cuneus gyrus | 10.4 | -91.6 | 14.7 | 120 | 303.74 | 0.000115 |
| Intraparietal sulcus | 26.9 | -57.7 | 47 | 22 | 10.81 | 0.003895 |
| **RSA: POST pcode "appearance" component (difference between zero-order and partial correlation)** | | | | | | |
| *Left hemisphere* | | | | | | |
| Calcarine sulcus, lingual gyrus, cuneus gyrus | -4.3 | -88.9 | 7.5 | 923 | 2440.4 | 1.73E-09 |
| Superior parietal gyrus | -21.7 | -65.8 | 53 | 166 | 249.2 | 0.001153 |
| Intraparietal sulcus | -23.6 | -62.6 | 33.3 | 77 | 128.47 | 0.000869 |
| Precentral sulcus | -49.5 | 0.5 | 33.6 | 28 | 56.47 | 0.000578 |
| *Right hemisphere* | | | | | | |
| Calcarine sulcus, cuneus gyrus | 6.6 | -90.4 | 15 | 1013 | 2832.4 | 1.55E-08 |
| **RSA: POST pcode "meanings" component (partial correlation with visual)** | | | | | | |
| *Left hemisphere* | | | | | | |
| The union of multiple sub-regions across occipital cortex and lateral occipito-temporal cortex | -12.4 | -85.6 | 6.8 | 2227 | 5099.55 | 8.23E-11 |

| | | | | | | |
|---|---|---|---|---|---|---|
| The union of multiple sub-regions including intraparietal cortex, temporo-parietal cortices, and posterior superior temporal cortex | -21.2 | -64.6 | 35.8 | 2012 | 3226.41 | 3.43E-07 |
| Precuneus | -9.7 | -55.2 | 35.3 | 367 | 623.27 | 8.16E-07 |
| Postcentral cortex | -34.3 | -36 | 45.6 | 403 | 607.18 | 9.79E-05 |
| Precentral cortex, inferior frontal sulcus | -46.5 | 4.6 | 17 | 338 | 632.33 | 4.34E-05 |
| Medial superior frontal gyrus | -6 | 4.3 | 61.5 | 198 | 543.47 | 7.72E-05 |
| Superior temporal sulcus | -47.9 | -36.9 | -4.3 | 171 | 241.56 | 0.000256 |
| Subcentral cortex | -55.3 | -19.1 | 18.7 | 83 | 175.26 | 0.001327 |
| Superior precentral sulcus | -36 | -6.9 | 45.3 | 43 | 97.11 | 0.000299 |
| *Right hemisphere* | | | | | | |
| The union of multiple sub-regions across medial occipital cortex, lateral occipito-temporal cortex, and temporo-parietal cortex | 8.8 | -87.3 | 6.5 | 2895 | 6707.16 | 2.43E-07 |
| Precuneus | 5 | -45.7 | 44.8 | 217 | 340.07 | 0.00019 |
| Intraparietal sulcus | 34.1 | -44.3 | 48.4 | 157 | 181.97 | 0.000183 |
| Inferior frontal gyrus | 47.2 | 23.3 | 6.8 | 74 | 136.66 | 4.15E-05 |
| **RSA: POST pcode "meanings" component (difference between zero-order and partial correlation)** | | | | | | |
| *Left hemisphere* | | | | | | |
| Calcarine sulcus, lingual gyrus, cuneus gyrus | -4.3 | -88.9 | 7.5 | 824 | 2195.84 | 2.47E-09 |
| Intraparietal sulcus | -23.6 | -62.6 | 33.3 | 57 | 94.41 | 0.002141 |
| *Right hemisphere* | | | | | | |
| Calcarine sulcus, lingual gyrus, cuneus gyrus | 6.6 | -90.4 | 15 | 901 | 2565.18 | 1.52E-08 |
| **Univariate code > pcode (POST)** | | | | | | |
| *Left hemisphere* | | | | | | |
| The union of multiple sub-regions including intraparietal cortex, angular gyrus, precuneus, lateral occipital cortex, and posterior inferior occipito-temporal cortex | -42.2 | -72.1 | -2.8 | 4234 | 7652.15 | 5.47E-10 |
| The union of multiple sub-regions in lateral frontal cortex | -45 | 38.1 | -5.7 | 2138 | 4594.39 | 1.96E-08 |
| Cingulate cortex | -1.3 | 10.7 | 25.7 | 455 | 743.8 | 7.35E-09 |
| Lingual sulcus, fusiform gyrus | -30.9 | -47.2 | -12.8 | 345 | 930.35 | 1.55E-07 |
| Medial superior frontal gyrus | -6.5 | 19.3 | 45.4 | 257 | 659.32 | 4.14E-08 |
| *Right hemisphere* | | | | | | |
| The union of multiple sub-regions including intraparietal cortex, angular gyrus, precuneus, lateral occipital cortex, and posterior inferior occipito-temporal cortex | 32.5 | -69.6 | 26.6 | 4531 | 9000.77 | 2.14E-12 |
| Inferior precentral sulcus, inferior frontal sulcus | 41.1 | 3.2 | 30.4 | 641 | 1213.73 | 4.37E-08 |
| Superior frontal sulcus, middle frontal gyrus | 24.1 | 2.7 | 48.4 | 438 | 816.41 | 2.24E-06 |
| Posterior cingulate cortex | 4.8 | -32.3 | 29 | 233 | 380.03 | 3.46E-06 |
| **Univariate pcode > code (POST)** | | | | | | |
| *Left hemisphere* | | | | | | |
| Calcarine sulcus, lingual gyrus, cuneus gyrus | -2.6 | -86.9 | -4.9 | 2014 | 5058.21 | 6.83E-14 |
| The union of multiple sub-regions including superior temporal cortex, temporal pole, supramarginal gyrus, subcentral cortex, and inferior insula sulcus | -53.7 | -10.1 | -11.5 | 2642 | 5345.99 | 3.18E-09 |
| Middle cingulate cortex, medial superior frontal gyrus | -6 | -6.4 | 57.5 | 755 | 1416.64 | 1.46E-06 |

| | | | | | | |
|---|---|---|---|---|---|---|
| Anterior/middle cingulate cortex | -8 | 44.2 | 1.8 | 579 | 1340.84 | 2.11E-05 |
| Postcentral sulcus | -28.5 | -37 | 53.9 | 222 | 343.42 | 6.67E-05 |
| Central sulcus | -39.4 | -21 | 42.4 | 191 | 403.21 | 0.00032 |
| Precentral gyrus | -43.3 | -12.1 | 50.1 | 128 | 323.16 | 0.000113 |
| *Right hemisphere* | | | | | | |
| Calcarine sulcus, lingual gyrus, cuneus gyrus | 8.2 | -79.6 | 10.9 | 1797 | 4631.86 | 1.21E-14 |
| Superior temporal cortex, temporal pole, middle temporal gyrus | 49.5 | -8.9 | -18 | 1329 | 3226.82 | 7.73E-10 |
| Anterior cingulate cortex, medial superior frontal gyrus, anterior middle frontal gyrus | 29.4 | 39.6 | 21.7 | 1166 | 2948.16 | 3.82E-07 |
| Supramarginal gyrus, posterior lateral sulcus | 53 | -30.8 | 29.3 | 930 | 1349.96 | 2.12E-07 |
| Anterior insula | 39.4 | 2.4 | -13.3 | 759 | 1417.82 | 6.57E-07 |
| Middle/posterior cingulate cortex | 6.2 | -23.5 | 41.6 | 365 | 644.63 | 2.98E-06 |
| Central sulcus | 35.2 | -18.8 | 38.9 | 270 | 482.95 | 3.68E-06 |
